# Ancient genomes illuminate Eastern Arabian population history and adaptation against malaria

**DOI:** 10.1101/2023.10.01.559299

**Authors:** Rui Martiniano, Marc Haber, Mohamed A. Almarri, Valeria Mattiangeli, Mirte C. M. Kuijpers, Berenice Chamel, Emily M. Breslin, Judith Littleton, Salman Almahari, Fatima Aloraifi, Daniel G. Bradley, Pierre Lombard, Richard Durbin

## Abstract

The harsh climate of Arabia has posed challenges in generating ancient DNA from the region, hindering the direct examination of ancient genomes for understanding the demographic processes that shaped Arabian populations. In this study, we report whole genome sequence data obtained from four Tylos-period individuals from Bahrain. Their genetic ancestry can be modelled as a mixture of sources from ancient Anatolia, Levant and Iran/Caucasus, with variation between individuals suggesting population heterogeneity in Bahrain before the onset of Islam. We identify the G6PD Mediterranean mutation associated with malaria-resistance in three out of four ancient Bahraini samples and estimate that it rose in frequency in Eastern Arabia from 5-6 kya onwards, around the time agriculture appeared in the region. Our study provides the first characterisation of the genetic composition of ancient Arabians, shedding light on the population history of Bahrain and demonstrating the feasibility of aDNA studies in the region.

## Introduction

Following flint and pottery evidence of human occupation from ∼5000 BCE^1^, from around 2200 BCE Bahrain can be connected with references in Mesopotamian cuneiform records to the Dilmun civilisation, both as a trading centre ^2^ and playing an important role in Sumerian mythology, including in their creation myth and in the Epic of Gilgamesh ^3,4^. This period also saw the start of a funerary tradition that ultimately resulted in the highest density of burial mounds in the ancient world (in the order of 100,000)^5^. The end of the Late Dilmun phase (∼600 BCE) coincided with the Persian Achaemenid conquest of Mesopotamia, whose influence in Bahrain (600-300 BCE) is attested by the presence of bowls and other artefacts typical from that culture ^6^.

Around 325 BC, an expedition to the Arabian coast sent by Alexander the Great reached the shores of Bahrain, which from that time onwards would be known as Tylos. The Tylos period (∼325 BCE - ∼600 CE) was characterised by exceptional prosperity and marked by Hellenistic and Persian influence ^7^. The death of Alexander and subsequent disintegration of the Macedonian Empire led to the establishment of the Seleucid Empire (312–63 BCE), which dominated a vast area comprising Anatolia, Levant, Mesopotamia and Iran, also controlling the eastern Arabian islands of Failaka and Bahrain. The Seleucids eventually lost authority in Mesopotamia, leading to the formation of the semi-independent Kingdom of Characene (141 BCE–222 CE) in Southern Iraq, a vassal to the Parthian Empire that would govern Bahrain until Sasanian conquest ^8,9^.

Recent aDNA studies from the northern regions of the Middle East revealed that the first farmers from Anatolia, the Levant and Iran each descended from local, genetically distinct hunter-gatherer populations, with variable amounts of gene-flow from neighbouring groups ^10,11^. For example, early Anatolians admixed with Mesopotamians and later with Levantine agriculturalists ^12^, whereas in the Levant, early farmers derived from Epipaleolithic Natufians and Anatolian Neolithic populations ^11^. During the transition from the Chalcolithic to the Bronze Age, admixture between various Middle Eastern groups intensified, leading to increasing genetic homogenisation in the region ^11^. Around this time, an ancient Iranian-related component was introduced into the Levant ^13,14^, followed by steppe/European-related ancestry in the Iron Age ^15,16^. In the neighbouring region of Mesopotamia, the ancient DNA record is still sparse, and therefore the genetic composition of the local hunter-gatherers remains unknown, but recently published Pre-Pottery Neolithic (PPN) genomes from Upper Mesopotamia are genetically intermediate along the ancestry cline extending from ancient groups from Anatolia/Levant to Iran/Caucasus ^12,17^.

Due to challenges associated with the recovery of ancient genomes from hot and humid climates^18^, no ancient genomes from the Arabian Peninsula have been published so far. Surveys of present-day genomes from Arabia and the Levant suggest that they were shaped by different demographic processes: first, Arabians carry an excess of East African- and Natufian-related ancestry in comparison with Levantine populations, who in contrast, bear higher proportions of European and Anatolian Neolithic ancestry ^19,20^. Second, Arabian and Levantine populations present different size trajectories, with the former being affected by a pronounced bottleneck event occurring around 6 kya that coincides with the ‘Dark Millenium’, a period of increasing aridification in Arabia ^21^, and the latter showing a more recent size reduction associated with the ∼4.2 kya climate event ^20^, of wider distribution across Anatolia, the Levant and Mesopotamia ^22^. Third, Arabians show increased levels of consanguinity in comparison to the Levant ^23,24^, potentially leading to higher prevalence of genetic disorders including G6PD deficiency ^25^.

To examine these topics directly using ancient genomes, we sequenced four individuals from the island of Bahrain dating from the Tylos period (∼300 BCE - 600 CE). Using modelling approaches, we determine that their ancestry is a mixture of sources from ancient Levant, Iran/Caucasus and Anatolia, with one individual presenting higher affinity to Levantine groups than the others, potentially as a result of historically recorded incursions of Arabian or Levantine tribes into Bahrain ^26^, and two individuals with additional CHG (Caucasus Hunter-Gatherer)/post-Neolithic Iranian ancestry, which can tentatively be explained by Iranian-associated influence in Bahrain during pre-Islamic times. Additionally, we detect the G6PD Mediterranean variant in three Tylos-period Bahrainis and estimate that it rose to high frequencies in Eastern Arabia due to strong positive selection exerted by malaria endemicity coinciding with the appearance of agriculture. Lastly, we detect large runs of homozygosity in one individual, suggesting consanguineous unions in pre-Islamic Tylos populations. The present work provides, for the first time, a snapshot of the genetic composition of ancient Eastern Arabia and demonstrates the feasibility of ancient DNA studies in the region.

## Results

### Ancient DNA sample sequencing and determination of authenticity

We extracted DNA from 25 skeletal samples from ancient burial mounds from the island of Bahrain ranging from the Dilmun period to the Tylos period and sequenced them to assess endogenous DNA preservation (Supplementary Text 1, Table S1). Of these, only four were sufficiently well preserved for additional sequencing, with the remaining samples presenting negligible amounts of human endogenous DNA. Here, we report shotgun sequence data obtained from these four samples, all derived from petrous bones, one from Abu Saiba (AS) and three from Madinat Hamad (MH1, MH2, MH3; Figure S1, Table 1, Supplementary Text 2). Of these samples, three were sequenced to approximately 1x, and one to 0.24x (Table 1). Due to poor collagen preservation, we could only obtain radiocarbon dates for two out of the four sequenced individuals, placing them in the Late Tylos/Sasanian period (‘LT’, ∼300-622 CE), with MH1 being older (432-561 cal. CE) than MH3 (577-647 cal. CE) (Figure S2). Sample MH2 was not directly dated, but its archaeological context places it in the Late Tylos period (Supplementary Text 2). The Abu Saiba sample was excavated from a cemetery with known occupation between 200 BCE-300 CE ^7,27^, and therefore it dates confidently within the boundaries of the Early/Middle Tylos Period (‘EMT’), more precisely during the times of Seleucid and Characene influence in Bahrain which preceded the emergence of the Sasanian Empire.

**Table 1.** - Ancient DNA samples from Bahrain sequenced in this study.

| SampleID | Dates | Reads | Coverage | Sex | Y-chromosome haplogroup | mtDNA haplogroup | mtDNA Contamination (%) | X-chromosome Contamination (%) |
| --- | --- | --- | --- | --- | --- | --- | --- | --- |
| AS_EMT | ~200 BCE-300 CE | 71,371,001 | 0.94 | XX | - | J1c15a1 | 1.36 | - |
| MH1_LT | 432-561 cal. CE<br>(1565 BP) | 69,192,646 | 1.06 | XY | J2a2a1a~ | R2 | 2.00 | 2.92 |
| MH2_LT | 300 - 600 CE | 80,197,765 | 1.26 | XX | - | T2b | 0.97 | - |
| MH3_LT | 577-647 cal. CE<br>(1456 BP) | 18,586,203 | 0.24 | XY | H2 | U8b1a2a | 0.64 | 2.00 |

Deamination patterns in these sequences conform with those typical of ancient DNA (Figure S3), supporting the presence of authentic ancient DNA sequences, and we observed low contamination rates on the mitochondria of all four samples (≤ 2%) and on the X-chromosome of the two male samples (< 3%, Table 1). None of the samples were close relatives of each other (Figure S4).

### Genome-wide affinities of Tylos-period individuals from Bahrain

To examine the genetic affinities of the four Tylos-period samples from Bahrain, we performed a principal component analysis (PCA) ^28,29^ on 1,301 present-day West Eurasians ^30,31^, including 117 recently reported Arabians and Levantines ^20,32^, on which we projected 529 ancient individuals (Figure 1A; full PCA shown in Figure S5). The four Bahrain Tylos samples were positioned intermediately along the cline connecting western (Anatolian and Levantine) and eastern sources (CHG and IRN_Ganj_Dareh_N) and in the vicinity of Upper Mesopotamian pre-pottery Neolithic individuals from Southeastern Turkey and Northern Iraq (Mesopotamia_PPN), Neolithic farmers from Armenia (ARM_Masis_Blur_N and ARM_Aknashen_N) and Azerbaijan (AZE_N). Also in the proximity of the ancient Bahrain samples are various post-Neolithic groups from Iran (IRN_HajjiFiruz_ChL/IRN_Hasanlu_IA, IRN_DinkhaTepe_A_BIA), Bronze Age groups from Eastern Turkey (TUR_Hatay_Alalakh_MLBA), Iraq (IRQ_Nemrik9_LBA), Syria (Syria_Ebla_EMBA) and various other Levantine groups (Hellenistic/Roman), all of which were previously reported to contain a mixture of two or more sources related to Anatolian, Levantine and Iranian Neolithic/CHG ancestries ^11–14,33,34^.

**Figure 1.**
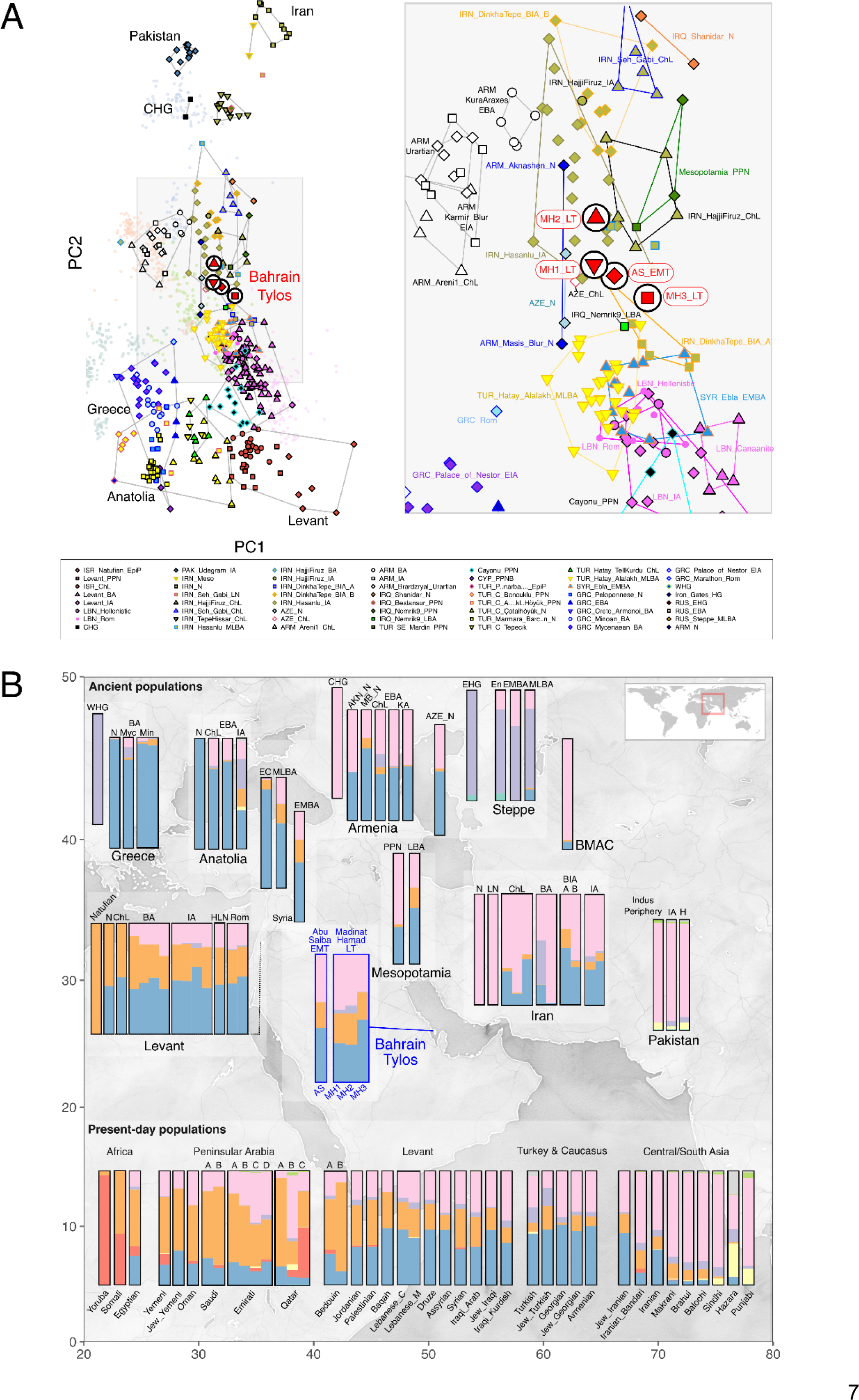
- A) Principal Component Analysis with 1,830 present-day and ancient Eurasians and 579,407 SNPs. Ancient samples are indicated with larger symbols as in the key. B) Pertinent results from a Dystruct analysis (K=9) of 2,073 samples, including 658 ancient samples binned into eleven different time periods, using 85,043 transversions and with the mean ancestral components estimated for each population. Ancient samples above and present-day samples below. Legend: AKN - Aknashen; BIA - Bronze Iron Age; BMAC - Bactria–Margiana Archaeological Complex; ChL - Chalcolithic; E/M/LBA - Early/Middle/Late Bronze Age; EC - Early Chalcolithic; H - Historical; HLN - Hellenistic; IA - Iron Age; KA - Kura-Araxes; MB - Masis Blur; Min - Minoan; Myc - Mycenaean; N - Neolithic; Rom - Roman.

The Bahrain_Tylos individuals are slightly differentiated. Compared with the older individual AS_EMT, samples MH1_LT and MH2_LT are shifted towards eastern populations from Armenia ChL-IA composed mainly of CHG- and Anatolian-related ancestry and variable amounts of steppe ancestry, with MH2_LT being further separated from the Bahrain group towards the direction of CHG and ancient groups from Iran and Pakistan. Sample MH3_LT is closer to ancient Levantine groups than the other three individuals from Bahrain.

To further investigate the ancestry composition of the Tylos individuals, we performed a temporally aware model-based clustering analysis on an expanded dataset using DyStruct (Figure 1B). At k=9, Anatolian_N, Natufians, WHG (Western Hunter-Gatherers), and Iran_N/CHG define independent ancestral components which contribute to the majority of ancient and present-day West Eurasians (Figure 1B; Figure S6). Consistent with the PCA results, the four Tylos Period individuals are broadly similar to and intermediate between post-Neolithic groups from the Levant (BA/IA/Hellenistic/Roman) and Iran (ChL/IA), albeit with higher amounts of Iran_N/CHG-related ancestry than ancient Levantines and higher Natufian-related ancestry than ancient Iranians. These results are corroborated by positive statistics for both f4(Mbuti, ISR_Natufians; Ancient Iranians, Bahrain_Tylos) and f4(Mbuti, IRN_Ganj_Dareh; Ancient Levantines, Bahrain_Tylos) which indicate higher allele sharing between Bahrain_Tylos and ISR_Natufian_EpiP in comparison with Ancient Iranians (Figure 2A), and between Bahrain_Tylos and IRN_Ganj_Dareh in comparison with Ancient Levantines (Figure 2B).

**Figure 2.**
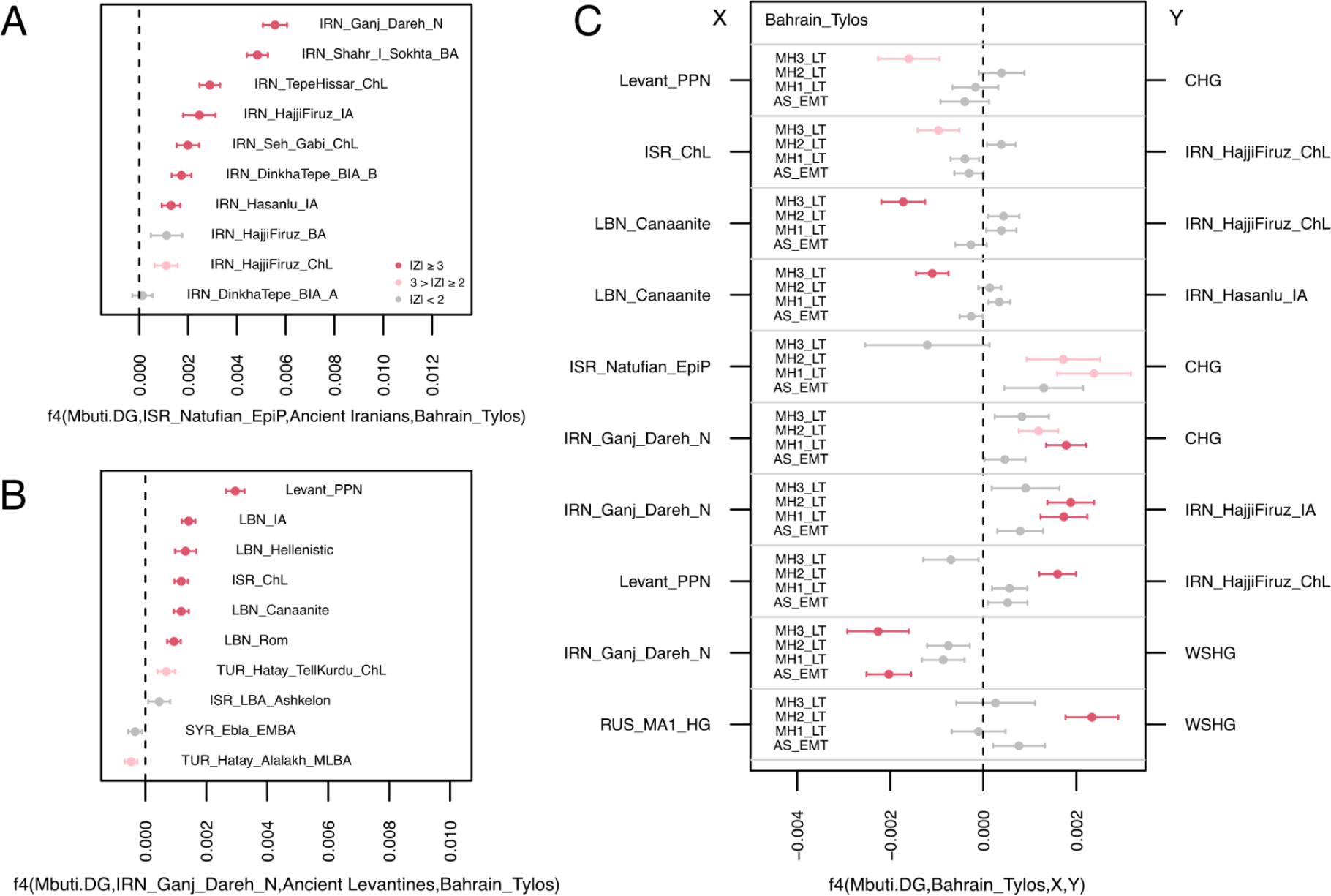
- Allele sharing statistics of the form A) f4(Mbuti, ISR_Natufians; Ancient Iranians, Bahrain_Tylos), B) f4(Mbuti, IRN_Ganj_Dareh; Ancient Levantines, Bahrain_Tylos) and C) f4(Mbuti, Bahrain_Tylos; X, Y).

We also observe some variability within the four Bahrain samples in the clustering analysis (Figure 1B), with MH3_LT carrying additional Anatolia_N and less Iran_N/CHG-related ancestry than the other samples. Here, Anatolian and Levantine ancestry were in some cases not well differentiated due to the presence of Anatolian ancestry in Neolithic Levantines. To investigate this, we estimate f4(Mbuti,X; MH3, AS_EMT) and confirm that sample MH3_LT presents excess allele sharing with both Anatolians (TUR_C_Çatalhöyük_N; Z=-2.11) and Levantines (LBN_Canaanite; Z=-2.06) in comparison with the earlier sample AS_EMT, and we obtain increased significance for both statistics (Z=-3.05 and Z=-3.79, respectively) when replacing AS_EMT with the Madinat Hamad Late Tylos pair MH1 and MH2 (Table S2), suggesting a gradient of Levantine-related ancestry which is maximised in MH3_LT and minimised in MH2_LT, with AS_EMT and MH1_LT presenting intermediate values between them, as reflected in the PCA (Figure 1A). In Figure 1B, it is also apparent MH1 and MH2_LT individuals present slight differences in comparison with the other two Bahraini samples, notably, higher Iran_N/CHG-related ancestry and an additional small amount of Western/Eastern Hunter Gatherer (WHG/EHG)-related ancestry (approx. 3.1-6.2%) which forms around half of the genetic composition of Steppe EMBA^35,36^ and can also be found in Armenia_ChL and Iranian HajjiFiruz_BA/IA.

To formally investigate these subtle differences, we tested for excess allele sharing between Bahrain_Tylos and ancient groups X and Y by estimating f4(Mbuti, Bahrain_Tylos; X, Y) (Figure 2C). We corroborate excess affinity between MH3_LT and Levantines, especially with LBN_Canaanites, in comparison with CHG or post-Neolithic Iranians, but we obtain no significant results for the remaining samples from Bahrain. Conversely, MH1 and MH2_LT present excess allele sharing with CHG in comparison with ISR_Natufians, suggesting lower Levantine and higher CHG ancestry in these samples, with AS_EMT presenting intermediate values. MH1 and MH2_LT also present increased CHG and IRN_HajjiFiruz_IA ancestry in comparison with IRN_Ganj_Dareh_N, as well as slightly higher West Siberian Hunter-gatherer (WSHG) ancestry particularly in MH2_LT.

### Modelling Bahrain Tylos ancestry

We found that Bahrain Tylos is best modelled as a mixture of sources from ancient Anatolia, Levant and Iran/Caucasus. First, we obtained significantly negative results (z≤-3) for the statistic f3(Bahrain_Tylos; X, Y) when X was an ancient Levantine or an Anatolian population (Natufians, Levantine or Anatolian Neolithic) and Y an ancient group from Caucasus or Iran (CHG or IRN_Ganj_Dareh_N) (Table S3), suggesting that these ancestries have plausibly contributed, even if distantly so, to the formation of Bahrain Tylos.

Second, two-way qpAdm models return many possible combinations, nearly always with sets of populations carrying three types of ancestry: Levantine, Anatolian and CHG/Iranian Neolithic (Table S4). For instance, all four samples can be modelled with mixtures of Levant_PPN and CHG (p>0.05; Figure 3A, Table S4). However, when replacing Levant_PPN with ISR_Natufian_EpiP, only two samples (AS_EMT and MH3_LT) were successfully modelled, although with lower p-values, suggesting that an additional Anatolian-related component, which is present in Neolithic Levant, but absent in Natufians and CHG, is required to model the remaining samples. We note that Anatolian Neolithic ancestry can also derive from other groups, such as ARM_Aknashen_N (formed of Anatolian and CHG ancestry), which returns feasible models when combined with ISR_Natufian_EpiP or Levant_PPN (p>0.05; Table S4).

**Figure 3.**
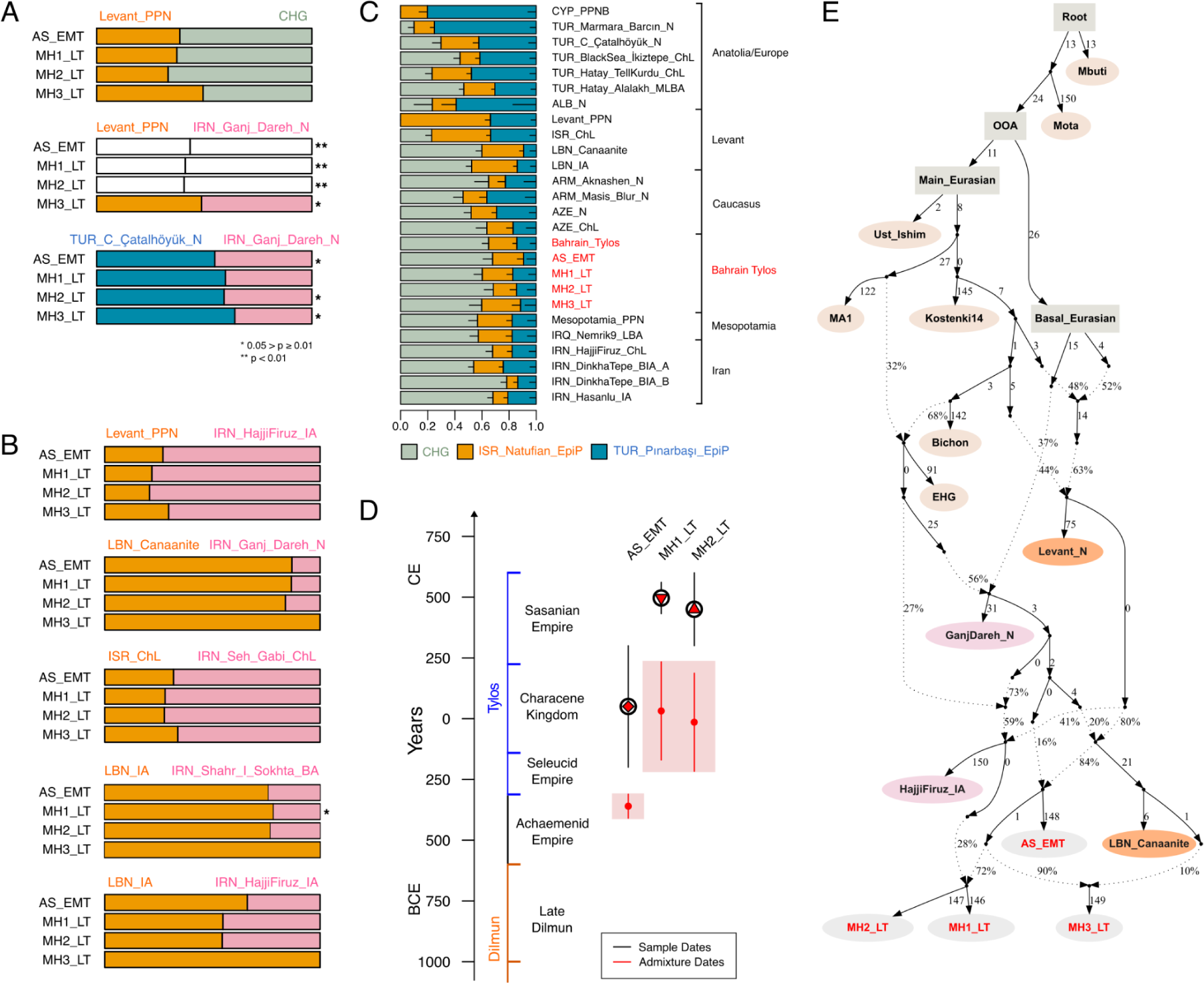
- Models of Bahrain Tylos ancestry. A) Selected rank 1 qpAdm models of the four Bahrain Tylos individuals including distal sources and B) a mixture of distal and proximal sources. Feasible rank 0 models are shown instead in the cases where the rank 1 model was rejected. * indicates 0.05 > p ≥ 0.01, ** and white bars indicate rejected models at p < 0.01. C) qpAdm models of ancient Near Easterners using Natufians, Pinarbasi and CHG as sources. D) Inferred time of admixture in Bahrain Tylos using DATES ancestry covariance decay curve using LBN_Canaanites and IRN_Shahr_I_Sokhta_BA as references representing ancient Levantine and ancient Iranian ancestries, respectively. E) Modelling Bahrain Tylos ancestry using qpGraph.

Models including Levant_PPN and IRN_GanjDareh_N sources are not as successful, given that only the lower coverage MH3_LT individual can be modelled in this way (p=0.042; Figure 3A; Table S5) suggesting that CHGs might be better representatives than Neolithic Iranians for the eastern ancestry present in the Tylos samples. Accordingly, we obtain a significantly negative (Z=-3.81) f4-statistic f4(Mbuti.DG, Bahrain_Tylos; CHG, IRN_Ganj_Dareh_N), suggesting excess affinity between Bahrain_Tylos and CHG in relation to IRN_Ganj_Dareh_N. Although, as seen in our dystruct analysis (k=12, Figure S6), CHG and IRN_Ganj_Dareh_N ancestries are difficult to distinguish from one another, and it is possible that they both contributed to the formation of Tylos-period Bahrainis. In fact, we also obtained generally less confident but feasible models when combining Iranian Neolithic or Late Neolithic with CYP_PPNB or TUR_C_Çatalhöyük_N, with the latter two populations containing both Anatolian and Levantine ancestry (Figure 3A, Table S4).

Third, we also find support for a tripartite genetic ancestry of Bahrain Tylos when including more temporally proximal groups as sources in our models (Figure 3B; Table S4), obtaining feasible combinations of older Levantine (Levant_PPN, ISR_ChL) and more recent Iranians (IRN_HajjiFiruz_IA), or older Iranians (IRN_Ganj_Dareh_N, IRN_Seh_Gabi_ChL) and more recent Levantines (LBN_Canaanite, LBN_IA), in addition to post-Neolithic sources from both regions. The overall pattern in these models is that the most recent source contributes with a greater amount of ancestry to Bahrain Tylos samples, irrespectively of being Iranian or Levantine, due to the fact that later samples tend to carry the Anatolia/Levant/Iran/CHG-related ancestries required for modelling the four samples from Bahrain.

Lastly, we evaluate the ancestry of Bahrain Tylos individuals in a context of Near Eastern variation by estimating ancestry proportions using a previously published ^12^ three-way model with TUR_Pınarbaşı_EpiP, ISR_Natufian_EpiP and CHG as sources in a set of relevant ancient groups which can also be modelled in this way (Figure 3C) ^12^. In this model, the Bahrain_Tylos samples present similar ancestry proportions to AZE_ChL, Mesopotamia_PPN and IRQ_Nemrik9_LBA, IRN_Dinkha_Tepe_A, LBN_IA and to ARM_Aknashen_N. Accordingly, rank=0 qpAdm models show that Tylos-period Bahrainis form a clade with several of these ancient groups (p≥0.01; Table S6), suggesting that similar sources have contributed to their ancestry.

### Recent gene-flow events shaped the genetic composition of Tylos-period Bahrainis

We next investigated the timing of admixture between ancient Levantine and Iranian sources in Bahrain Tylos samples using DATES (Figure 3D). We observe different admixture times for AS_EMT and MH1-2_LT: the first occurred 14±1.7 generations before AS_EMT (357-455 years, assuming a generation time of 29 years) coinciding with a period of Achaemenid influence in Bahrain, and the second occurred 16±7 generations (261-667 years) before MH1_LT and MH2_LT, a period defined by the incorporation of Bahrain into the Characene Kingdom, a vassal state to the Parthians.

We summarise these processes in an admixture graph (Figure 3E). In this graph the earliest sample from Abu Saiba (AS_EMT) derives from admixture between sources related to the Levant and Iran_N/CHG. The three Late Tylos individuals descend from admixture between the earliest sample AS_EMT and other sources, with MH1 and MH2_LT having an additional pulse of post-Neolithic Iranian-related ancestry, here represented by IRN_Hajji_Firuz_IA, whereas the MH3_LT sample received additional ancestry associated with the Levant, represented by LBN_Canaanite from Sidon. These inferences are supported by the observation of additional IRN_IA ancestry in MH1 and MH2_LT relative to AS_EMT and MH3_LT, and that MH3_LT bears more Levantine ancestry than the other samples from Bahrain (Figure 3B). We show an alternative model where the post-Neolithic Iranian source is instead represented by IRN_Shahr-i-Sokhta in Figure S7.

### Affinities with Mesopotamia

Considering the abundance of evidence of contacts between Ancient Bahrain and Mesopotamia, we evaluated separate models using Mesopotamia_PPN (two Pre-pottery Neolithic samples from Northern Iraq and one from Mardin in Turkey), a Late Bronze Age (LBA) sample from Iraq, and the Upper Mesopotamian Cayonu_PPN from Southeastern Turkey as a source. Firstly, we observe that all four Tylos samples form a clade with IRQ_Nemrik9_LBA, two samples form a clade with Mesopotamia_PPN (MH1 and MH3_LT) and only the lower coverage MH3_LT forms a clade with Cayonu_PPN (Table S6). When exploring two-source models with Mesopotamia_PPN, we can model all samples with Levant_PPN as a second source, except for sample MH2_LT, which requires additional EHG ancestry, or using RUS_AfontovaGora3, SRB_Iron_Gates_HG or LBN_IA as a second source instead of Levant_PPN (Table S7). While we cannot reject various models, the low Z-scores attributed to the admixture weights for most models, especially for those corresponding to non-Mesopotamian sources, do not allow for establishing the precise role of Mesopotamian groups in the formation of Bahrain Tylos.

### Tylos-Period Bahrainis are genetically closer to present-day Levantine populations than to present-day Arabians

Regarding affinities with present-day populations, the temporally aware model-based clustering analysis (Figure 1B) suggests that Tylos period Bahrain samples are more similar to present-day Levantine groups than to present-day Arabians or South Asians, who show higher and lower amounts of Natufian ancestry, respectively. To investigate this in more detail, we performed a haplotype-based ChromoPainter and FineSTRUCTURE analysis with a dataset including Tylos-period Bahrainis, ancient samples from the Levant ^13,37^ and Iran ^10^ with available whole genome shotgun sequence data, as well as present-day individuals from the Human Origins dataset ^1138^, and from Arabia and the Levant ^20^ (Figure 4).

**Figure 4.**
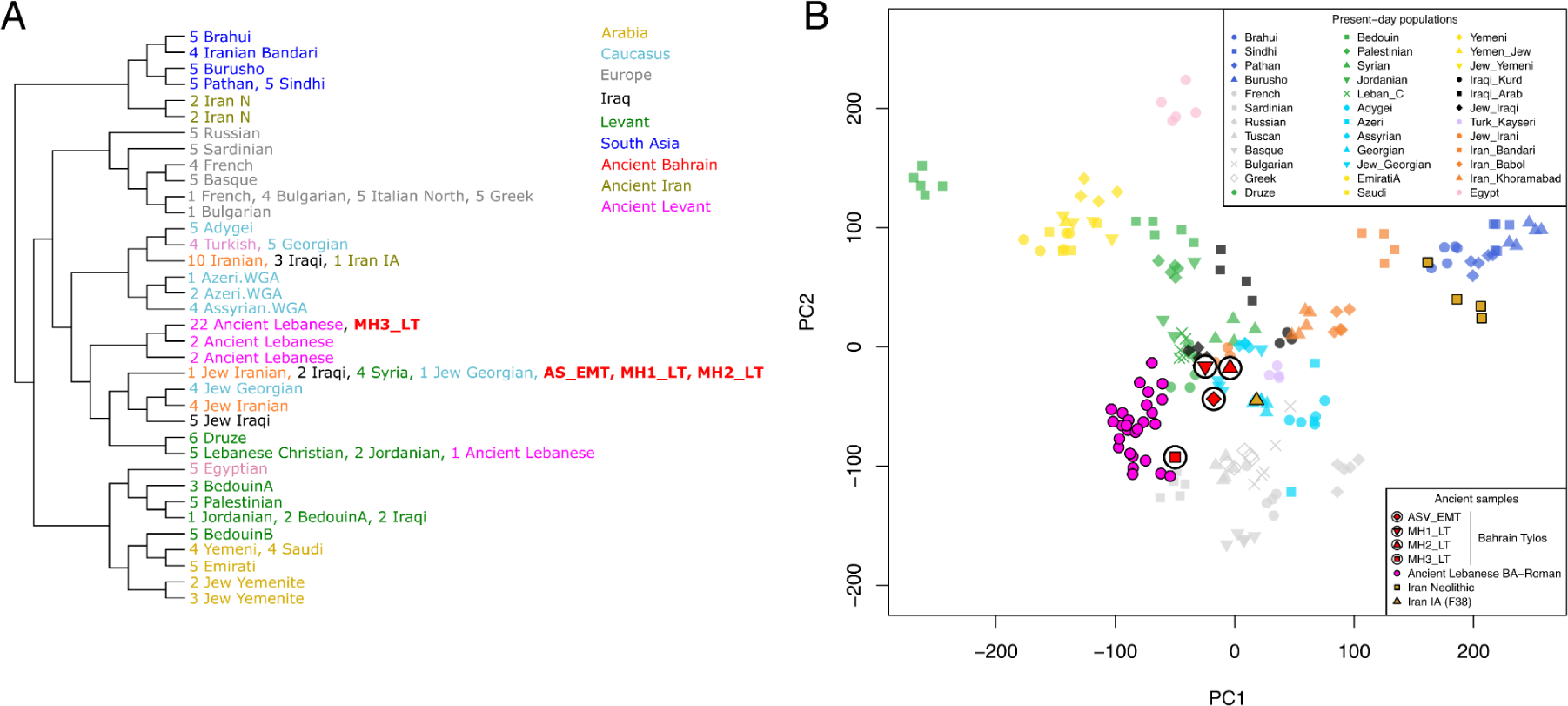
- A) FineSTRUCTURE phylogeny and clustering based on haplotype sharing patterns from present-day Eurasians and ancient Middle Easterners . B) PCA estimated from the ChromoPainter haplotype sharing matrix.

Three Bahrain Tylos samples (AS_EMT, MH1 and MH2_LT) were included in a cluster composed of Syrians, Iraqis and two Jewish individuals from Iran and Georgia (Figure 4A), whereas sample MH3_LT clustered with 22 ancient individuals from Lebanon ranging from the Bronze Age to the Roman Period. In a PCA of the haplotype sharing matrix (Figure 4B), we observe that MH3_LT is closer to ancient Lebanese and Sardinians, corroborating our previous inferences of increased ancient Levantine and Anatolian Neolithic ancestry in this sample. The remaining three Bahrain Tylos individuals are positioned closer to present-day groups from the Levant, Iraq and the Caucasus, including several Jewish populations from these regions, and between ancient Lebanese and an Iranian IA individual.

In order to gain further insights into the relationship of Tylos to present-day populations from the Arabian Peninsula and the Levant, we tested if the Bahraini samples form a clade with any modern population in our dataset using a set of reference populations (as outgroups) that can differentiate the different ancestries in the Near East. We found that Iraqis, Assyrians and Jewish groups from Iran, Georgia and Iraq could derive all their ancestries from Tylos-period Bahrainis (Table S8). Arabians such as Saudis, Emiratis and Yemenis have, in addition to ancestry from Tylos-period Bahrain, ancestry from East Africa, while Levantines such as Druze and Lebanese have additional Southeast European ancestry (Table S9). Here we should note that modelling present-day populations with Tylos-period Bahrain does not imply a direct contribution but rather that they are suitable representatives of the ancestry found in present-day populations.

### Phenotypic prediction

We used HIrisPlex-S ^39^ to predict hair, skin and eye colour phenotypes of the Bahrain_Tylos samples. All four samples were predicted to have brown eyes (>99% probability). For hair colour, the prediction was either Brown (∼50%) or Black (∼50%). Two samples (MH2_LT and MH3_LT) were predicted to have “Dark” skin pigmentation (>90%), whereas the results for the remaining two samples were less certain, with AS_EMT potentially having relatively lighter skin pigmentation (Table S10). The predicted phenotypes found in ancient Bahrain are similar to present-day Middle Easterners and South Asians. Within these regions, subtle geographical trends occur at the level of skin pigmentation. Specifically, non-Bedouin Levantine groups from the HGDP tend to have higher proportions of “Intermediate” skin pigmentation, while Pakistanis tend to have higher proportions of “Dark” skin than most Levantine populations, especially in the South ^39^.

### Diverse Uniparental Lineages link Ancient Bahrainis to Middle Eastern populations

We used pathPhynder ^40^ to examine the Y-chromosome lineages of two male samples from the Late Tylos-period Bahrain in a context of present-day and ancient samples (Figure S8, Table S11). Individual MH3_LT carried the H2 haplogroup which is associated with the spread of Near Eastern and Anatolian farmers into Europe ^41^. MH3_LT was placed in the same clade as a PPNB Jordanian and two Anatolian samples (Alalakh_MLBA and Ilipinar_ChL), which is consistent with the genetic affinities of this individual (Figure S8A).

Sample MH1_LT, the only other male sample in our dataset, presented the J2a2a1a∼ lineage, which in a phylogeny with 2,014 individuals is carried by a present-day Brahmin and an Uighur individual. In the aDNA record, this lineage and derived haplotypes have been identified in two Turkish individuals (Gordion_Anc and Mediaeval), in a Canaanite and in an Iron Age Hasanlu individual ^12,42^, and in various present-day Central Asian samples from Turkmenistan and Kazakhstan. An eastern origin (Iran-Caucasus region) for haplogroup J2 is likely, given that the earliest occurrence of this lineage in the aDNA record is in hunter-gatherers from Caucasus and Iran ^11,43^, with the latter being placed at the base of the J2a clade (Figure S8B). MH1_LT’s inclusion in this Y-chromosomal clade is consistent with the subtle excess in shared autosomal ancestry with CHG/IRN_HajjiFiruz_IA individuals in this sample (Figure 2C).

Strikingly, neither of these lineages were found in present-day Arabians ^20^. Their presence in both western (Anatolia/Levant) and eastern (Iran/Caucasus/Central Asia) ancient samples reflects connectivity between these ancient civilisations as early as the pre-pottery Neolithic (as attested by the presence of eastern ancestry and J2 Y-chromosome lineage in Cayonu_PPN ^44^) which apparently intensified during post-Neolithic times ^42^.

The mitochondrial lineages of the Tylos-period samples suggest maternal ancestry sharing with various groups from the Near East, Caucasus and South Asia. The earliest sample AS_EMT carries the mtDNA J1c15a1 haplogroup previously reported in two present-day samples from Iraq ^45^ and Azerbaijan ^46^, groups who typically carry Iranian- and CHG-related ancestry. The Late Tylos samples from Madinat Hamad presented distinct lineages, with MH1_LT belonging to the R2 haplogroup which is predominantly South Asian (southern Pakistan and India), but also distributed in the Middle East, Caucasus and Central Asia^47^. In the ancient DNA record, this lineage is most frequent in Iranian groups, including three Neolithic and two BIA/LBA individuals ^42,48^, which is consistent with the additional post-Neolithic Iranian-related ancestry in this sample. Interestingly, its presence in a Bronze Age Canaanite, coinciding with the emergence of Iranian ancestry in the Levant, provides additional support for the association of R2 lineages with this source of ancestry. Sample MH2_LT carries the T2b lineage, of widespread distribution in European samples. Lastly, MH3_LT presents the U8b1a2a haplogroup, also found in two ancient LBA Armenians, two Turkish (one ChL and the other dated to 750-480 BCE) ^42^ and in a present-day Jordanian individual ^49^, suggesting distribution in the Caucasus, Levant and Anatolia, and consistent with the increased Levantine and Anatolian affinities of this sample. These observations support extensive female as well as male migration in ancient West Asia including Eastern Arabia.

### Runs of Homozygosity suggest a spectrum of inbreeding similar to present-day Middle Easterners

We also examined runs of homozygosity (ROH) in the Tylos samples and compared them with present-day and ancient genomes (Figure 5A). Contemporary Middle Eastern populations show long tracts of ROHs which are indicative of recent consanguinity, while ancient regional hunter-gatherer groups are known to have relatively large numbers of shorter ROH, reflecting their smaller population size^50^. We observed an excess of ROH size in sample MH2_LT which is similar to that seen in present-day Middle Eastern and South/Central Asian populations (Figure 5A). In comparison to ancient groups, we find that three individuals (AS_EMT, MH1 and MH3_LT), show a similar ROH distribution to Bronze and Iron Age regional populations (Figure S9). However, one sample (MH2_LT) appears to have larger ROHs, with one longer than 16Mb, suggesting direct parental relatedness (Figure S10).

**Figure 5.**
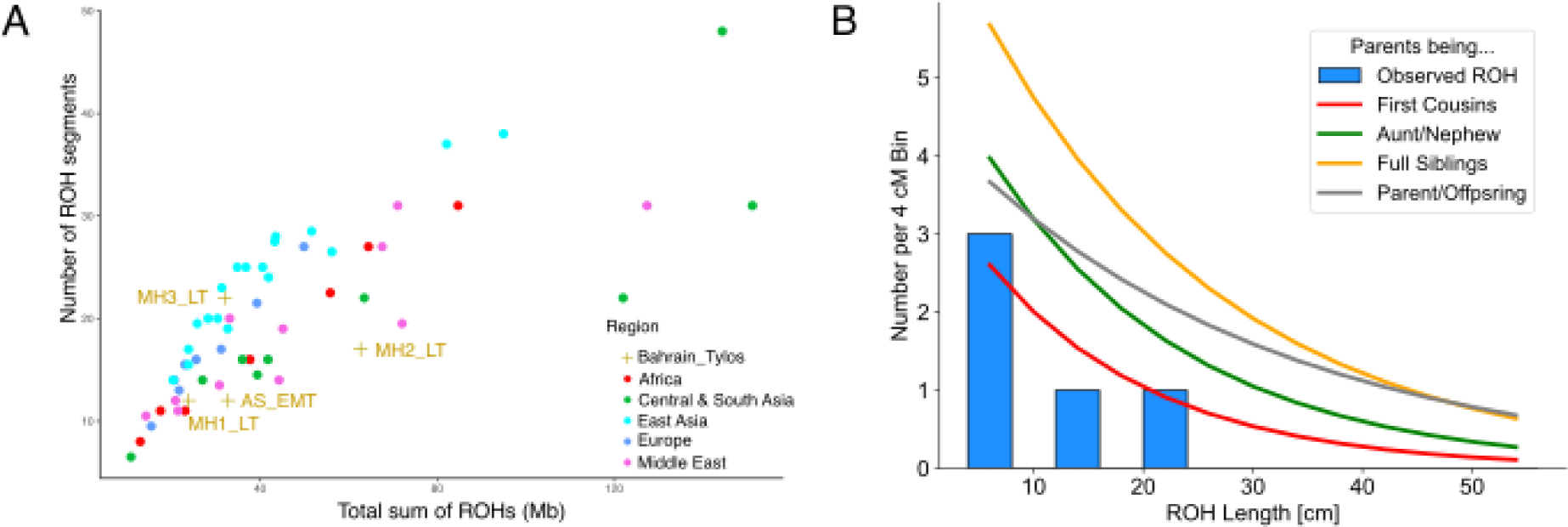
- A) Runs of homozygosity in Bahrain_Tylos and worldwide populations (plotting median values for each population). B) ROH Length (>4cM) distribution in sample MH2_LT and expected ROH distribution for the offspring of different consanguineous unions.

To confirm this result, we then used an alternative method^51^ for identifying ROHs in ancient samples. We observe that MH2_LT has levels of inbreeding consistent with being the offspring of related individuals (potentially second cousins; Figure 5B; Figure S11A), including a stretch on chr7 that spans 20.47 cM (Figure S11B). This finding is evidence that consanguineous union was likely to be already practised in pre-Islamic Arabian societies.

### High prevalence of the malaria-protective G6PD Mediterranean mutation in ancient Eastern Arabia arising with the introduction of agriculture

G6PD deficiency is the most common enzymatic defect in humans ^52^ and its distribution in worldwide populations correlates with regions currently or historically affected by malaria, including Africa, the Mediterranean, the Middle East and Southeast Asia ^53^, leading to the proposal of a link between G6PD deficiency and malaria protection ^54^.

Considering that malaria was endemic in Bahrain during historical times ^55^, with osteological analyses suggesting that it was present in the island at least by the Tylos period ^56^, we examined the Tylos-period Bahraini samples for the presence of mutations putatively-associated with malaria protection. Two ancient samples from Bahrain carried the G6PD Mediterranean mutation (rs5030868; G>A; p.S188F). MH2_LT is potentially heterozygous, given that we observed two reads overlapping this SNP in this sample, one supporting the derived allele A and the other with the ancestral allele G, whereas AS_EMT presented two reads with the derived allele.

To corroborate our findings and obtain additional insights about the distribution of this variant in ancient populations, we imputed the X-chromosome of the Bahrain_Tylos samples together with 37 ancient Lebanese and four ancient Iranians. We confirm the presence of derived alleles (genotype probability > 0.95) at the rs5030868 locus in samples AS_EMT (homozygous), MH2_LT (heterozygous) and MH3_LT (male; hemizygous mutant). None of the ancient Lebanese we imputed carried this mutation and neither did five individuals from Bronze Age Greece with sequencing reads spanning this site ^57^ but we observed a heterozygous genotype in a Western Iranian Neolithic sample (AH1, genotype probability > 0.9). When estimating a phylogeny with the imputed X-chromosome haplotypes containing the mutation of interest, we observe the inclusion of three Tylos-period Bahrainis in a clade with additional samples carrying the derived allele, including present-day Emiratis, Omanis and Yemenis, as well as the Iranian Neolithic sample AH1 (Figure 6A).

**Figure 6.**
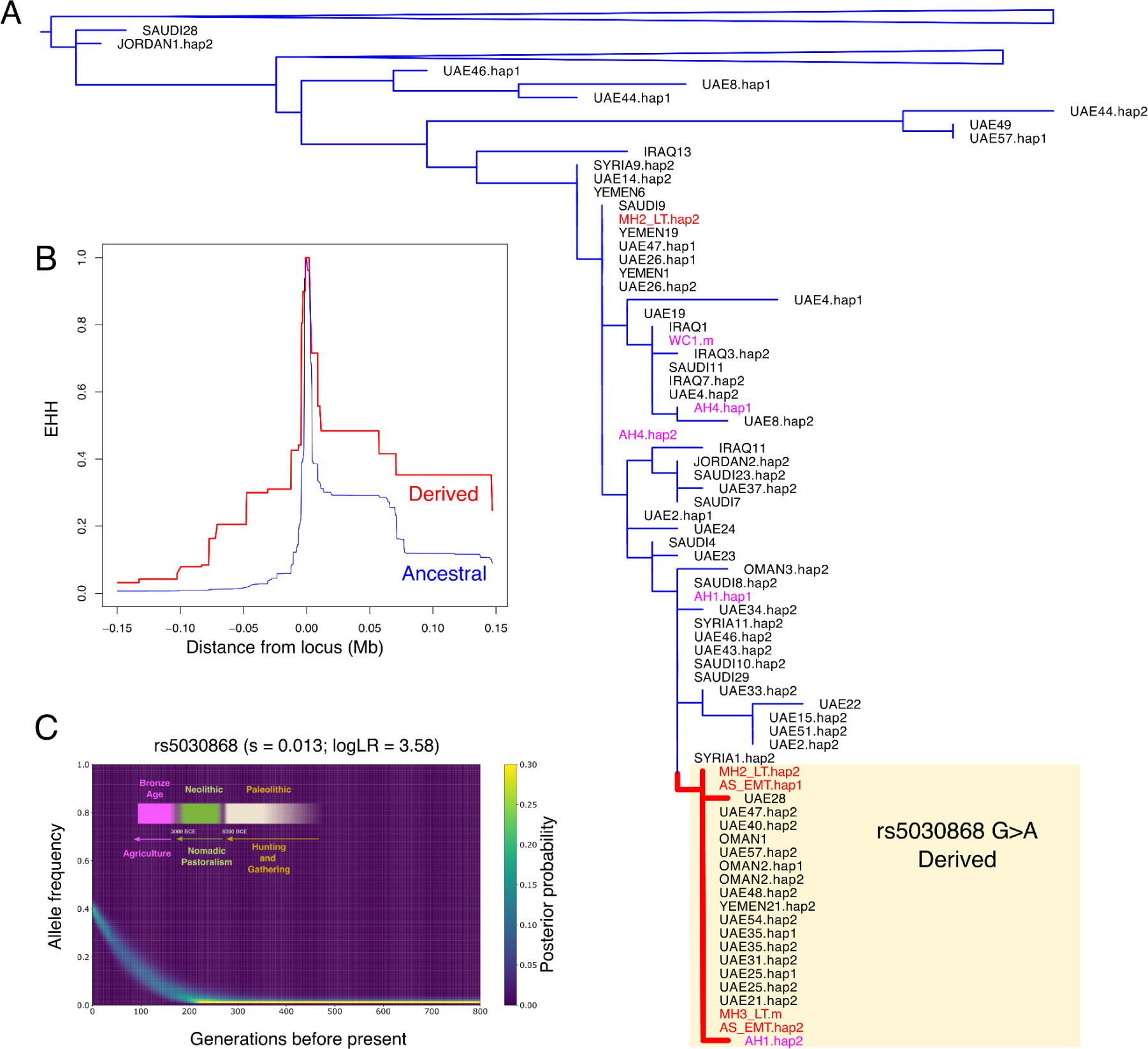
- A) Phylogeny of Eastern Arabian and Levantine haplotypes surrounding the G6PD Mediterranean mutation (rs5030868). B) Extended Haplotype Homozygosity in present-day Arabians. C) Allele frequency trajectory of the variant in EmiratiC highlighting changes in modes of subsistence in Eastern Arabia.

In present-day populations, we observe relatively high frequencies of this variant in Makrani (0.19), Brahui (0.14) and Pathan (0.08) population samples, all of which bear high levels of Iranian Neolithic-related ancestry. In Europe frequencies are substantially lower, with a single Sardinian sample carrying the derived allele in the HGDP dataset (0.03), and three out of 53,000 European individuals in the gnomAD database ^58^. In the Middle East, it can be found in Palestinians (0.06), but not in Druze, Bedouins, Jordanians or Iraqis ^20^. In Arabian populations, the derived allele reaches its highest frequencies in EmiratiC (0.38), a subgroup of the Emirati who carry substantial Iranian/South Asian-related ancestry (Figure 1B), and 6% in the rest of the Emirati population. It has also been found in the present-day populations of Yemen (0.04), and Qatar (0.03)^32^, but not in Saudis ^20^. In the Levant and Eastern Arabia, the Mediterranean mutation is responsible for the majority of G6PD-deficient cases, reaching very high frequencies in Bahrainis (0.91)^59^, Northern Iraqi (0.88), Kuwaitis (0.74) and Jordanians (0.54) ^60^.

In order to search for signatures of selection associated with the G6PD Mediterranean, we performed an extended haplotype homozygosity analysis (EHH) on the haplotypes surrounding this SNP in a dataset of Arabian and Levantine groups. We observe that the haplotypes carrying the derived allele tend to be longer than those with the ancestral allele (Figure 6B), consistent with a signature of positive selection. We subsequently modelled the historical change in frequency of the variant and found that it has increased rapidly in the past six thousand years in EmiratiC with an estimated selection coefficient of 0.013, suggesting strong selective pressure (Figure 6C). This date broadly coincides with the start of the Bronze Age in Eastern Arabia (∼3200 BCE), a period of cultural transformation marked by the shift from nomadic pastoralism to agriculture ^61^.

Overall, these findings suggest that the G6PD Mediterranean variant may have already been present, albeit at low frequencies, in Iranian Neolithic samples, and was introduced into the Levant and Arabia through admixture, rising to high frequencies in Eastern Arabians after the appearance of agriculture.

## Discussion

### Population heterogeneity and admixture in Tylos-period Bahrain

In the present work, we address an important gap in the ancient DNA record by presenting the first whole-genome sequences from Eastern Arabia, more precisely from Tylos-period Bahrain, an era of Hellenistic, Parthian and Sassanian influence in the region. We observe that the four individuals from the Tylos period occupy an intermediate position along the genetic ancestry cline spanning from western (Anatolian/Levantine) to eastern (Iran/CHG) sources, with some degree of heterogeneity in ancestry composition within our samples.

We detected an excess of Levantine-related ancestry in sample MH3_LT, which raises the possibility of admixture with populations from the Levant or as yet unsampled Arabian groups that may have carried this ancestry into Bahrain before or around the onset of Islam. The first hypothesis is supported by the substantial affinity of MH3_LT to Lebanese Canaanites and other Levantine groups, and by shared Y-chromosomal and mitochondrial DNA lineages between this individual and ancient eastern Anatolians and Levantines. The second hypothesis is supported by the existence of historical records attesting the presence of several nomadic Arab tribes, such as the Abd Al-Qays and Al-Adz, under Nasrid control ^62^ in Bahrain during this time. These tribes migrated from Southwestern Arabia into Bahrain and neighbouring regions in pre-Islamic times, subsequently participating in the conquests of Persia and Mesopotamia in the second half of the 7th century CE^26^. Such migrations could have brought additional Levantine ancestry to Bahrain, given that Southwestern Arabians, such as the Yemeni from Maarib, carry substantial Natufian-related ancestry but virtually no African ancestry ^19^, as is the case of sample MH3_LT, and are, therefore, potentially good representatives of Arabian ancestry prior to admixture with other sources including Africans.

The presence of minor steppe-related ancestry in the Late Tylos samples (MH1 and MH2_LT), but not in the earliest sample from Abu Saiba (AS_EMT), does not require a direct influx from Steppe groups, associated, for example, with the LBA migrations that spread Indo-European languages into South Asia ^33,63^. Instead, this ancestry may have been introduced to Bahrain through admixture with geographically and temporally more proximal groups from ancient Levant or Iran where it appears around the Iron Age ^15,37^ and Bronze Age ^33^, respectively. Considering that Bahrain history is marked by Achaemenid, Parthian and Sasanian influence, it is plausible that groups associated with these empires acted as vectors for introducing steppe ancestry into Bahrain. In support of this hypothesis, the J2a2a1 Y-chromosome lineage of sample MH1_LT is also present in the Iranian IA and in other ancient groups with substantial Iranian-related ancestry (C. Asian, Anatolian, Levantine BA), but neither of the two males reported here carried the R1a-Z93 lineage associated with the spread of Steppe ancestry and Indo-European languages into South Asia ^33^.

We inferred admixture times between Levantine and Iranian sources occurring first, during the Achaemenid Empire, and second, during a time of Characene rule of Bahrain. Interestingly, there was no substantial overlap between our admixture time estimates and the period of Seleucid influence in Bahrain, suggesting limited gene-flow during this period of Greek cultural influence. However, this analysis suffers from several limitations. First, it is possible that this finding may derive from a lack of power to determine exact admixture times in the case of continuous gene-flow. Second, the lack of direct radiocarbon dates for two of our samples, AS_EMT and MH2_LT, limits precise estimation of admixture timing. Third, we currently lack Achaemenid Period genomes for comparison to determine with more certainty the extent of genetic changes at the time of the Seleucid Empire.

Given the limited sample size (n=4) and the narrow temporal focus on a specific period in Bahrain’s history, it is important to emphasise that it remains challenging to establish whether these ancestry differences derive from specific population movements occurring alongside political changes in the Near East after the Dilmun Period or if they reflect pre-existing genetic diversity within the island’s population. As seen in our PCA (Figure 1A), Near Eastern groups such as Iron Age Levantines and Iranians present relatively high diversity, and therefore, it is feasible that the differences we observe between our samples derive from preexisting genetic structure within the Bahraini population not directly related to the events mentioned here. Furthermore, while we were able to identify several plausible models for describing the ancestry of Tylos period Bahrainis, the current lack of ancient genomes from the Arabian Peninsula and sparse sampling from Mesopotamia (particularly from the south, from where no aDNA samples have been published so far) prevents us from testing models with more proximal sources relevant to Bahrain. Nevertheless, various analyses point to some degree of genetic similarity between a Late Bronze Age sample (IRQ_Nemrik9_LBA) and the Bahraini samples, which gathers support from archaeological and textual evidence of contacts between Mesopotamia and Bahrain at the time of the Dilmun civilisation. However, this observation derives from a single Mesopotamian sample, and therefore this connection should be reexamined when additional ancient genomes become available. A particularly important question remains regarding the ancestry of the Dilmun-period inhabitants, which, once characterised, will help describe more precisely how the genetic composition of Arabians has evolved through time, the relationship of Dilmun with neighbouring civilisations, and their contribution to later Tylos-period groups.

### Increased Malaria endemicity in Eastern Arabia

We report the presence of the G6PD Mediterranean mutation in three out of four samples and four out of six alleles from ancient Bahrain. Notably, this includes a homozygous female and a hemizygous male, who would have suffered G6PD deficiency and possibly hemolytic anaemia. This finding suggests that this mutation occurred at appreciable frequencies in Eastern Arabia during the Tylos period. The identification of this variant in an Iranian Neolithic sample, but not in ancient Levant or ancient Europe, together with its prevalence in present-day groups from Pakistan and in Arabian groups with high Iran N-related ancestry (EmiratiC), suggests a West or South Asian origin for this variant rather than European/Mediterranean, contradicting the hypothesis that it was introduced into the Middle East in relatively recent times alongside the Greek expansions of the 1st millennium BC ^64^ or that its dispersion into Europe occurred alongside Neolithic migrations ^65^. Our data suggests instead that this mutation may have been disseminated into Arabia through admixture with Mesopotamian/Iranian/S. Asian groups, rising to high frequency due to selective pressures, particularly along the Arabian coast, where malaria incidence was especially high. According to our estimates, the onset of selection for this variant occurred between 5 and 6 kya, broadly coinciding with the emergence of agriculture in Eastern Arabia. Such changes are evidenced by the presence of domesticated cereals at Al-Hili dated to around 3000 BCE ^61^ and the development of oasis agriculture associated with the Umm an-Nar culture (2700-2000 BCE) ^66^. Increased sedentarism in areas with available water ^67^ would have created a propitious environment for the proliferation of malaria, providing selection pressure for malaria-protective mutations.

Our confidence in imputing the Mediterranean variant in the three samples that carried the derived allele is supported by direct observation of the variant in a total of three reads spanning that position in two individuals. Although this SNP is a G/A change, potentially causing difficulties in distinguishing it from post-mortem deamination, this is likely to be more problematic for the Iranian Neolithic sample than for our UDG-treated samples. Nevertheless, this Iranian individual shares the same haplotype as the other derived samples, providing additional confidence in it being a carrier. Expanding our investigation of the Mediterranean variant to a wider set of ancient groups would be desirable, but currently challenging because this SNP is not included as part of the 1240k SNP target capture array commonly used for sequencing ancient samples ^68^. Future studies generating additional whole genomes from the Middle East and South Asia should provide more insights about the spread and distribution of malaria-protective variants in the ancient world.

Lastly, we demonstrate the feasibility of ancient DNA studies in Arabia, paving the way for future research aimed at elucidating human population movements in the region. The sequence data and insights here reported will be of long term importance for studying human population history in the Middle East and beyond.

## Methods

### Archaeological sample processing, DNA extraction, library preparation and sequencing

We collected 25 human remains from the Bahrain National Museum and the Qal’at al-Bahrain Museum belonging to the Dilmun and Tylos periods. We excised the denser part of the petrous portion of the temporal bones or in case of teeth, the root, which we converted to powder using a homogenizer (Retsch Mixer Mill). Where possible, sample homogenization was done progressively by shaking the bone multiple times for approximately 10-15 seconds, keeping the resulting powder between each session. This process allows the selection of the densest part of the petrous bone. We extracted approximately ∼130 mg of bone powder per sample as previously described ^69^ with modifications ^70^ and prepared double-stranded DNA libraries using a previously published method ^71^ with the modifications described in ^72^ (Supplementary Text 1). We amplified DNA libraries and performed initial sequencing runs on an Illumina MiSeq to assess endogenous human content and deamination patterns in the DNA libraries. Based on initial screening results, we prioritised the four most promising libraries for additional sequencing on a HiSeq X.

Radiocarbon dating of the four samples that were successfully sequenced was performed at Oxford Radiocarbon Accelerator Unit with fragments of the petrous portion of the temporal bone (Supplementary Text 2).

### Sequence read processing and alignment

We trimmed adapter sequences from high-throughput sequencing reads using AdapterRemoval v2 ^73^ with the parameters (--interleaved-input --minlength 30 –trimns --trimqualities --minquality 2 --collapse), and aligned these to the human reference genome (GRCh38, obtained from ftp://ftp.1000genomes.ebi.ac.uk/vol1/ftp/technical/reference/GRCh38_reference_genome/G RCh38_full_analysis_set_plus_decoy_hla.fa) using bwa aln ^74^, with the parameters -n 0.01 -o 2 and selected reads with mapping quality ≥ 25 using samtools ^75^, as previously recommended ^76,77^ and removed sequence read duplicates using sambamba markdup ^78^. We used the same steps to process published shotgun WGS ancient DNA from the Levant ^13,15,37^ and Iran ^10^.

### Sex determination and relatedness analysis

We determined the sex of the four ancient samples from Bahrain using a previously published method^79^ (Table 1). We performed a kinship analysis on the four ancient samples from Bahrain using READ (https://bitbucket.org/tguenther/read/) ^80^ with default parameters on approximately 1,240k SNPs of the Allen Ancient DNA Resource, genotyped as described below, which did not reveal any close kinship relationships between them.

### Authenticity of aDNA sequences and contamination estimates

To evaluate aDNA sequence authenticity, we estimated post-mortem deamination patterns in aligned sequence reads using PMDtools v. 0.6 (https://github.com/pontussk/PMDtools) ^81^. X-chromosome contamination in males was estimated using ANGSD ^82,83^ and mitochondrial DNA contamination for all four samples was estimated using contamMix v. 1.0-10 ^84^ using a panel of 311 present-day mtDNA sequences ^85^.

### Population Genetics Analyses

#### Variant calling

For population genetics analyses, we used the following datasets: the Allen Ancient DNA Resource (AADR; Allen Ancient DNA Resource version 44.3); present-day Middle Easterners ^20,32^, present-day worldwide populations from the Human Genome Diversity project (HGDP) ^38^ and from the Simons Genome Diversity Project ^86^ and previously _published ancient individuals_ ^1^_0–17,30,33–35,37,42,43,63,72,87–11_^2^ Where necessary, we converted the coordinates of the published data to the human genome assembly GRCh38 using CrossMapp v0.6.4 ^113^.

For the Bahrain_Tylos samples, we trimmed 2bp at the end of each read using bamUtil (https://github.com/statgen/bamUtil), while for the published Levantine and Iranian WGS we trimmed 3bp. We subsequently called variants at each position of the 1240k SNPs included in AADR using samtools mpileup, disabling base quality score recalibration and imposing a minimum base quality filter of q20. We generated pseudo-haploid genotypes by randomly sampling one allele at each SNP site using pileupCaller (https://github.com/stschiff/sequenceTools).

We genotyped present-day Middle Easterners ^20,32^ using bcftools v1.9 ^114,115^ with command “bcftools mpileup -T -q30 -Q30 | bcftools call -c” on the 1240k positions and merged the datasets using the mergeit program from the EIGENSOFT package v7.2.1^28,29^ with options docheck: YES and strandcheck: YES filtering sex-linked and triallelilic SNPs and sites that were outside the genomic accessibility mask ^38^, leaving 1,099,436 SNPs in the final dataset.

#### Principal component analysis and model-based clustering

We ran Principal Components Analysis (PCA) using smartpca v18140 from the EIGENSOFT package with parameters numoutlieriter: 0, lsqproject: YES, autoshrink: YES, projecting ancient samples on the principal components estimated with present-day Eurasian populations.

We also ran DyStruct (https://github.com/tyjo/dystruct) ^116^ from K=7 to K=12 in a dataset of 2,073 samples and 85,043 transversions with default parameters across eleven time points binned around the following times (generations ago): 450, 350, 250, 180, 150, 120, 90, 70, 50, 25, and present-day.

#### F-statistics, qpWave, qpAdm and qpGraph

We estimated f3- and f4-statistics and qpAdm using the AdmixTools 2 R package ^117^. For modelling the ancestry of Bahrain Tylos samples with qpAdm using ancient populations as sources, we used the following right populations: Mbuti.DG, CHG, EHG3, ISR_Natufian_EpiP, MAR_Taforalt_EpiP, RUS_AfontovaGora3, SRB_Iron_Gates_HG, TUR_Pınarbaşı_EpiP and WHG4. For qpWave tests using present-day populations, we used qpWave v1520 from the AdmixTools v7.0.2 package (https://github.com/DReichLab/AdmixTools) ^31^ with option allsnps:YES to test if ancient Bahrain formed a clade with any other ancient or modern populations. We used qpAdm v1520 to estimate ancestry proportions in present-day populations, using the following reference population samples: Mbuti, Ust’-Ishim, Kostenki14, MA1, China_Tianyuan, WHG, WSHG, Natufian, Levant_N, Morocco_Iberomaurusian, Anatolia_N, Iran_N, CHG, EHG, and Steppe_Eneolithic. We used qpDstat v980 also from the ADMIXTOOLS package to estimate D- and f-statistics to test for admixture and differentiation in ancient samples. We generated admixture graphs using qpGraph v7580 with parameters lsqmode:YES and inbreed:NO.

#### Chromopainter analysis

We ran the ChromoPainter/finestructure v4.1.0 inference pipeline^118^ with default settings outputting a co-ancestry matrix with values depicting haplotype segments shared between individuals. We ran the pipeline on a merged dataset composed of the 41 imputed ancient samples and 168 Modern samples from the Middle East, Europe and South Asia ^11,20^. We used 448K SNPs which have a minor allele frequency >5% in the reference panel and excluded sites with missingness more than 5%, and jointly phased the merged dataset using the 1000G high coverage reference panel^119^ using Beagle v5.2 ^120^.

#### Imputation

As Middle Eastern populations are not found in the commonly used 1000 Genomes Imputation panel ^121^, and some variants of interest are found in the Middle East but are very rare or even absent in the 1000G panel ^20^, in this study we created a reference panel which includes samples from the HGDP whole-genome sequencing study ^38^ and present-day Middle Easterners ^20^. The HGDP has several regionally-relevant populations from the Levant and South Asia, while Almarri et al. (2021) sampled populations from Arabia, the Levant and Iraq, and additionally included samples with present-day Iranian and South Asian ancestry. A total of 1064 samples were used to construct the reference panel, 929 from the HGDP and 135 samples from Almarri et al. 2021.

We first jointly called SNVs identified from both studies, excluding singletons, doubletons, and multiallelic variants, using GraphTyper_v2.6.2 ^122^ and set variants with GQ < 20 to missing. We limited the analysis to the strict genome accessibility mask ^38^, which covers ∼73% of the genome. As a reference panel cannot have missing data, we used Beagle 4.1^123^ with the option gtgl, which uses the genotype likelihood at sites set as missing, to identify the most likely genotype using the variation within the panel. We subsequently used a stringent filter by only including high quality sites (AR2 >0.97). The resulting reference panel comprised a total of 18.2 million SNVs.

We then used Beagle 5.1 ^124^ to phase the reference panel using default parameters. As the samples in Almarri et al., 2021 were physically-phased using linked-read sequencing, we were able to compare the samples (statistically-phased in our reference panel, with physically-phased from Almarri et al., 2021) to evaluate phasing accuracy of the panel. Extracting variants on chromosome 1 (840,284 SNVs), we used vcftools v0.1.16 with option --diff-switch-error to calculate the switch error at heterozygous sites per sample. We find a relatively low average switch error across all samples (mean 0.44%, stdev 0.1%), although we note the switch error rate will decrease by removing singletons and doubletons.

We subsequently imputed diploid genotypes of ancient samples using GLIMPSE v1.1.0 ^125^, using default parameters with the exception of increasing the number of main iteration steps of the algorithm to 15. We first called genotype likelihoods for each sample at sites found in the reference panel using the command “*bcftools mpileup -f {reference_genome} -I -E -a ’FORMAT/DP’ -T {variants_in_reference_panel} {sample}.bam -Ou | bcftools call -Aim -C alleles*”. To assess imputation accuracy, while taking into account our mapping and filtering process, we downsampled chromosome 21 of a published regionally-relevant early Neolithic genome (WC1)^10^ sampled from the Zagros region of present-day Iran from ∼10x to 0.1x, 0.25x, 0.5x, 1x and 2x using samtools view -s option. Genotype consistency was measured using the GLIMPSE_concordance tool by comparing the imputed variants with the variants called using all the read information (∼10x coverage) using only confidently called sites that have a minimum depth of 8 and minimum genotype probability of 0.999, a total of 85,848 SNVs. We find that for variants in the reference panel with minor allele frequency higher than 5% in the reference panel, even at 0.1x depth variants can be imputed with relatively high accuracy (r2 = 0.76), which increases with higher depth (0.25x - r2 = 0.89, 0.5x - r2 = 0.95, 1x - r2 = 0.97, 2x - r2 = 0.98). Subsequently we imputed the ancient Bahraini samples (4 samples) using the steps above, in addition to 37 published ancient Levantine and Iranian whole-genome sequences ^10,13,37^. The lowest coverage of the 41 imputed ancient samples was 0.24x.

To further improve the imputation quality, we followed a two-step imputation approach as previously described ^126^. We set to missing any variant that had an imputed GP < 0.99, and then re-imputed these missing variants using the larger high-coverage 1000G panel as a reference ^119^ using Beagle v5.1 ^124^. To further check the quality of the imputed genotypes, we compared Principal Component Analysis (PCA) and ADMIXTURE results between the imputed diploid and pseudohaploid calls across common variants (MAF > 5%). For both analyses we merged our ancient samples with the HGDP. In the PCA, we projected both the imputed and pseudohaploid calls on principal components calculated using modern samples (Figure S12). For ADMIXTURE, we ran the merged dataset using different Ks, from 3 to 8 (Figure S13). We find similar results between the pseudohaploid calls and the imputed calls, suggesting that the imputation performed is of relatively high quality.

For rs5030868, we created a special imputation panel solely of Middle Eastern samples (9% frequency) as rs5030868 is very rare in the 1000G panel (0.08%). We used GLIMPSE as stated above to impute common SNPs (>5%) covering a 3Mb region on the X chromosome surrounding the variant (5151 SNPs). We created a phylogeny of the locus using SNPs ±50Kb around rs5030868 with Seaview v5.0.5 (PhyML - HKY85 model, default conditions)1_2_7.

#### Relate

To estimate the allele frequency trajectory and selection coefficient of rs5030868 we ran RELATE v1.1.7 ^128^ to create a sequence of genealogies across 3Mb surrounding the variant as stated above using 59 female samples from the Almarri et al., 2021 Middle East dataset setting -m 1.11e-8 and -N 20,000. We subsequently extracted the EmiratiC individuals from the tree and estimated their population size history using EstimatePopulationSize.sh and then ran 100 samples using SampleBranchLengths.sh to sample branch lengths of rs5030868 from the posterior using a mutation rate of 1.11e-8 and supplying the .coal file from the previous step. We then ran CLUES ^129^ (inference.py script) with the option –coal to account for population size history.

#### Extended Haplotype Homozygosity

We conducted an EHH analysis on the region surrounding SNP rs5030868 using selscan v1.2.0 ^130^ with parameter --keep-low-freq and --ehh-win 5000.

#### Runs of Homozygosity

We identified ROHs using PLINK v1.9 ^131^ on the diploid imputed ancient genomes and present-day samples from the WGS HGDP ^38^ and Arabians and Levantines ^20^ using 652K common SNPs that were linkage-disequilibrium pruned (--indep-pairwise 50 5 0.9), and filtered for allele frequency (MAF > 5%) and missingness (< 5%). We also ran hapROH v0.64 to identify large ROHs (>4cm) and estimate parental relatedness^51^.

#### Uniparental lineage determination

We used pathPhynder ^40^ for investigating Y-chromosome lineages in the two male ancient samples from Bahrain and mtDNA-Server ^132^ for determining mitochondrial haplogroups.

#### Phenotypic Traits

We used the diploid imputed genotypes to explore phenotypic traits (eye colour, hair colour and skin colour) predicted by the HIrisPlex-S software (^133^ https://hirisplex.erasmusmc.nl/). Out of the 41 variants used by the software, we provided 38 variants and set the remaining 3 as “NA”. Two of these variants are very rare in frequency, while one was ambiguous when flipping and correcting for strand. We checked the quality of the imputation for the 38 variants, and found that the imputation INFO score generated by GLIMPSE for these variants was high (average INFO score 0.98, minimum 0.96), suggesting they were imputed with relatively high accuracy.

#### Time of admixture

We used DATES v4010 (Distribution of Ancestry Tracts of Evolutionary Signals) ^33^ to estimate the time of admixture in the Tylos samples. We set binsize: 0.001, maxdis: 1.0, runmode: 1 and used as source populations the ancient Bronze/Iron Age Levantines and Iranians: LBN_Canaanite, LBN_IA, ISR_Canaanite_MLBA, JOR_LBA, SYR_Ebla_EMBA, IRN_DinkhaTepe_BIA_A, IRN_DinkhaTepe_BIA_B, IRN_Hasanlu_IA, IRN_Shahr_I_Sokhta_BA.

#### Data Availability

Raw and processed sequence reads are made available at the European Nucleotide Archive, accessions XXXX and XXXX.

Imputed variants are made available in VCF format at XXXX. [Data will be made available upon publication]

#### Code Availability

All code used for analysis has been previously published and is publicly available.

## Acknowledgements

R.M. was funded by Wellcome grant 207492 and National Geographic Early Career Grant EC-61437R-19; R.D. by Wellcome grant 207492. The Abu Saiba excavation was funded by the French Ministry of Foreign Affairs. D.G.B was supported by Science Foundation Ireland/Health Research Board/Wellcome Trust Biomedical Research Partnership Investigator award no. 205072, ‘‘Ancient Genomics and the Atlantic Burden’’. M.A.A. was supported by Dubai Police GHQ. We thank the Wellcome Sanger Institute DNA Pipelines core for sequencing. We thank the President of Bahrain Authority for Culture and Antiquities, Shaikh Khalifa Bin Ahmed Al-Khalifa, for supporting the present research project and for providing access to archaeological samples for ancient DNA analysis. For the purpose of open access, the author has applied a Creative Commons Attribution (CC BY) licence to any Author Accepted Manuscript version arising from this submission.

## Author contributions

R.M. and F.A. conceived the project; B.C., S.A. and P.L. selected and provided archaeological samples; P.L. provided information about the archaeological context with contributions from B.C. and J.L.; R.M. processed bone samples; V.M. and E.B. carried out extractions and prepared sequencing libraries; R.M., M.H., M.A.A. and M.K. carried out analyses and interpreted them; R.M., M.H., M.A.A., P.L. and R.D. drafted the manuscript, and all authors commented on and approved the manuscript.

