## Supplementary Materials for "Ancient genomes illuminate Eastern Arabian population history and adaptation against malaria"

**Affiliations:**
1 - School of Biological and Environmental Sciences, Liverpool John Moores University, Liverpool, United Kingdom
2 - Institute of Cancer and Genomic Sciences, University of Birmingham Dubai, United Arab Emirates
3 - Department of Forensic Science and Criminology, Dubai Police GHQ, Dubai, United Arab Emirates
4 - College of Medicine, Mohammed Bin Rashid University of Medicine and Health Sciences, Dubai, United Arab Emirates
5 - Smurfit Institute of Genetics, Trinity College Dublin, Dublin 2, Ireland
6 - Department of Ecology, Behavior and Evolution; School of Biological Sciences; University of California San Diego; La Jolla; CA; USA
7 - Archéorient UMR 5133, CNRS, Université Lyon 2, Maison de l'Orient et de la Méditerranée - Jean Pouilloux, Lyon, France
8 - School of Social Sciences, University of Auckland, Auckland, New Zealand
9 - Bahrain Authority for Culture and Antiquities, Manama, Kingdom of Bahrain
10 - Mersey and West Lancashire Teaching Hospitals NHS Trust, Whiston Hospital, Warrington Road, Prescot, L35 5DR, Liverpool, United Kingdom.
11 - Department of Genetics, University of Cambridge, Cambridge, United Kingdom

[#] Lead contact
* Corresponding authors
[§] These authors contributed equally to this work.

### Supplementary Text 1 - Sample processing and DNA sequencing

##### *DNA extraction*

We sampled 25 petrous bones and teeth from ancient Bahrain individuals from 5 archaeological sites: Jari al-Shaikh (n=2), Shakhourah (n=3), Madinat Hamad (n=6), Saar (n=6) and Abu Saiba (n=8) (Table S1). Sample processing was carried out in dedicated ancient DNA facilities at Trinity College Dublin, Ireland, where all the precautions for ancient DNA processing were followed as described in [1]. We exposed all skeletal material to UV light for 15 minutes on either side to remove surface contaminants, and we cleaned the outer layer of bone with a drill before extraction. We targeted the densest section of the otic capsule region of the petrous temporal bone, which we pulverised for DNA extraction with a mixer mill. We used a subset of ~0.13 g of the bone powder for the following extraction step.

We extracted 25 samples using a silica column method [2] with 2 initial washing steps of 0.5M EDTA solution (EDTA1). We performed a second extraction for four samples selected from the results of the initial screening based on higher endogenous content and deamination was present. This time, we extracted both the second EDTA wash (EDTA2) and a new subset of the same bone powder treated with an initial washing step by 0.5% bleach solution (BEX1) [3].

The samples that were sequenced to high coverage were extracted as follows:

- MH1_LT - BEX1
- MH2_LT - EDTA2
- MH3_LT - BEX1
- AS_EMT - BEX1

##### *Library preparation and sequencing*

We performed an initial screening of each sample by constructing a double-stranded DNA NGS library using the method outlined in [4], with modifications as in [5]. Every library was first screened on a MiSeq Illumina platform (50 bp SE) at Trinseq (Ireland). DNA extracts selected for high-coverage sequencing were incubated with Uracil-DNA-glycosylase (UDG) enzyme (volume of 5 µl to 16.50 µl of extract) for 3 h at 37 °C to repair post-mortem molecular damage prior to a second library construction with the former method.

In order to increase complexity when sequencing for high-coverage, several PCRs with unique indexes were prepared from each library. Different PCR cycles were used according to the template concentration. We measured DNA concentration and assessed library quality using the Agilent Tapestation 2200 system with a D1000 screentape. High-coverage sequencing for all the selected libraries was carried out on a HiSeq X platform at the Wellcome Sanger Institute.

### Supplementary Text 2 - Archaeological sites and sample description

#### **2.1 ABU SAIBA**, Kingdom of Bahrain

*26°12'57.5" N  50°30'01.2" E*

Excavation : *French Archaeological Mission in Bahrain*, from 2017, under the direction of P. Lombard and J. Cuny (bio-anthropologist : B. Chamel).

The Abu Saiba necropolis, located 8 km west of Manama, the capital of Bahrain, appears as a low tell about 70 m in diameter and 4 to 5 metres high. During road works in 1983, its southern and western fringes were preventively explored by the Bahrain Directorate of Archaeology & Museums under the direction of Kh. Al Sindi and J. Littleton.

Since 2017, the French Archaeological Mission in Bahrain regularly excavates this typical necropolis from the Tylos period. Up to now, about 101 graves have been identified, and 90 excavated and studied according to the methods of archaeo-thanatology recommended by H. Duday [6]. These graves were individual burials in masonry pits of good architectural quality, several of which had yielded well-preserved furniture allowing to date them between the 1st century BC and the 1st century AD, a chronological frame confirmed by a set of calibrated radiocarbon dates. The cemetery, however, was extensively looted, presumably shortly after the burials; archaeological and anthropological traces are visible on the field, and their detailed study sometimes allows the reconstruction of the chronology of the actions of the looters [7,8]. The Abu Saiba necropolis displays a very homogeneous funerary architecture. Except for the perinatal and foetus, generally buried in pottery jars, all burials took place in masonry pit tombs of variable size depending on the deceased's age. The dimensions vary from 2.70 × 0.72 m. for a 1.20m. depth for larger ones to 0.96 × 0.42 m. for children's graves. The walls of these very regular rectangular pits are lined with rubble covered, as well as the bottom of the chamber, with a thick layer of very resistant lime mortar. At the top, a kind of bench, carefully coated, is arranged to receive the covering slabs, themselves sealed with mortar. The whole burial was generally covered with a mound of sandy sediment. The specific arrangement of a layer of small limestone slabs above several of them is usually considered a marker of high status. It likely indicates the burial of a particular group member [7]. The tombs are densely packed, the covering tumuli overlapping each other and gradually merging into a single low mound.

In the graves, the deceased's position is systematically the same, with the body placed on its back and its arms alongside it. Wood remains, or plank fragments in several graves, indicate the frequent use of wooden coffins. When the tomb was not affected by looting, it generally yielded a rich set of funerary offerings, including ceramics, alabaster or glass containers, frequent jewellery pieces and other adornments, cosmetic instruments, and objects of daily life. The placing of a small silver coin (obol) in the deceased's mouth (to pay the ferryman to the afterworld) confirms the strong links between Tylos and the neighbouring Hellenistic cultures of the Middle East [7].

One individual from Abu Saiba produced genome-wide data and is included in genetic analyses.
- AS_EMT (Grave 11, 2018) is from a heavily perturbed burial, which yielded very slender bones, in a good state of preservation, from an adult of indeterminate sex. A large amount of hypoplasia on the teeth indicates biological stress during childhood (starvation or childhood illness).

#### **2.2 MADINAT HAMAD (HAMAD TOWN)**, Kingdom of Bahrain

26°5'38.6"N / 50°30'5.85"E = site DS3

Excavation : *Bahrain Directorate of Archaeology and Museums*, 1981-1991.

In the archaeological literature, the generic name of "Madinat Hamad" refers to several archaeological sites, still present or now removed for urbanistic development. Located 18 km southwest of Manama, the capital of Bahrain, this vast sector is covered by a new city developed in 1984 under the impetus of the then-crown prince, who gave it its name. The area of Madinat Hamad, which extends from North to South over almost 9.5 km, was the subject of systematic preventive excavation campaigns between 1981 and 1991.

The area of Madinat Hamad is best known for its many Bronze Age burial mound fields associated with the Dilmun culture, and mostly dated between 2050 and 1750 BC [9–11]. Several thousand graves have been excavated and studied at approximately forty preventive excavation spots. The still unexcavated sectors of the three main Bronze Age cemeteries in this area (Buri, Karzakkan and Dar Kulayb), including 6,347 burial mounds, are today preserved and listed as World Heritage by UNESCO since 2019 under the respective names of Madinat Hamad 1, 2 and 3. They best represent the unique funerary landscape generated by the Dilmun phase necropolises in Bahrain island. Almost all of the Bronze Age burials studied in Madinat Hamad belong to the so-called "Late Type", the most frequently used during the apogee period of Dilmun, as opposed to the "Early Type", more commonly built at the end of the 3rd millennium in other areas of Bahrain Island [11,12]. This "Late Type" points to individual graves composed of a rectangular funerary chamber built on the ground in dry stone and covered with capstone slabs. Such a chamber was placed in the centre of a ring wall which initially gave these tombs built above ground the shape of small "towers". Following their slow collapse and steady erosion, they progressively took on their current appearance of rounded stony tumuli. Only one body, with few exceptions, was placed in each grave chamber. The skeleton is always found in a semi-flexed position on its right side, and generally oriented toward the north or the northeast, with hands brought close to the face. The body seems to have been directly deposited to the ground and was not covered with earth. It was regularly accompanied by burial offerings consisting of everyday or decorative ceramic artefacts, stone and copper vessels, copper weapons and personal ornaments (generally modest jewellery sets, semi-precious stone beads, necklaces and stone seals). These artefacts reflect the local productions of Bahrain island or witness the intensive trade between Dilmun and Mesopotamia, Iran, the Oman peninsula and the Indus Valley. Bones of ovicaprids, chicken or fish (remains of a funeral meal?) are sometimes part of the offering. It is difficult to determine the exact content of the funeral furniture due to the frequent, if not systematic looting of these graves during Antiquity.

The Madinat Hamad area was also reoccupied in later phases. One of the preventively excavated sectors, labelled "DS3", yielded between 1984 and 1986 nearly twenty small necropolises attributed to the culture of Tylos, whose identified remains essentially cover the period from c. 200 BC to c. 500 AD [13,14]. Each necropolis covered by a single mound seems to correspond to a specific social group and may include several dozen graves, or even more, positioned concentrically around a so-called "foundation" grave, where an eminent member (possibly the founder) of the group was buried. Most of the Tylos mounds of the Madinat Hamad Area DS3 developed in a sector previously occupied by Early Dilmun burial mounds. In a few cases, one observes that several Tylos graves directly reused former Dilmun burial chambers. However, most of the Tylos burials belonged to the traditional type, i.e. a perfect rectangular pit grave (average size: length:2m, width 0.54m, depth 0,72 m) with inside walls made of rough stones held together by mortar and covered with a layer of plaster which extended around the top to form a ledge to the closing capstones. A small mound of earth was originally covering each grave progressively merged with the neighbouring successive ones, producing the low mound covering the whole cemetery[15].

In 50 percent of tombs, the DS3 area revealed single burials with a skeleton laid on its back, hands on hips or alongside the body. In the remaining tombs, two or more successive primary burials were found in the same grave, separated by short or extended periods. Such a practice of multiple interments was most frequently observed in child burials[13]. The frequency of multiple burials is very high at DS3 compared to the usual Tylos burial practices in Bahrain and could point to a longer chronology for these graves. Despite these burials' frequent and heavy antique looting, funeral offerings appear to be almost systematic and quite diversified. They originally consisted of various containers in pottery, metal, stone or glass associated with personal items or adornments. The deposit of these artefacts reflects a whole system of belief (notably in a necessary journey to the afterlife) and echoes a very particular set of customs and traditions [15].

The DNA samples from Madinat Hamad Sector DS3 come from Mounds 30 and 70a, excavated in 1985-86. Mound 30, roughly round, was fully excavated. It measured c. 30 x 30m and had a maximum height of 2.58 m above the surrounding ground and revealed 41 graves of Tylos type built above five dating from the Dilmun phase. Mound 70a was of smaller size and was not excavated in totality. It yielded 27 graves of Tylos type [14].

Three individuals from Madinat Hamad produced genome-wide data and are included in genetic analyses.

- MH1_LT was excavated from Mound 30 at DS3, grave 28, which contained a minimum of four commingled burials. Ancient DNA was extracted from a petrous bone from an undetermined individual. Radiocarbon dated: 432-561 cal AD (1565 BP)
- MH2_LT was excavated from Mound 70a at DS3, grave 1, which contained a minimum of eight commingled burials. Ancient DNA was extracted from a petrous bone from an undetermined individual. Grave finds were attributed to Phase III (50-150 AD), however, the tomb was used multiple times [14].

- MH3_LT was excavated from an undetermined location at DS3. Ancient DNA was extracted from a petrous bone from an undetermined individual. Radiocarbon dated: 577-647 cal. AD (1456 BP).

#### 2.3 Radiocarbon dating

Radiocarbon dating was performed at the Oxford Radiocarbon Accelerator Unit (ORAU) and resulted in the following dates: MH1_LT (BAH_MTF) 1565±20 BP, 432-561 cal. AD (OxA-38603); MH3_LT (BAH_N) 1456±22 BP, 577-647 cal. AD (OxA-38708). It was not possible to obtain radiocarbon dates for MH2_LT (BAH_MHM) and AS_EMT (BAH_ASV) due to insufficient collagen yield in the bone samples provided for analysis.

#### Supplementary Figures

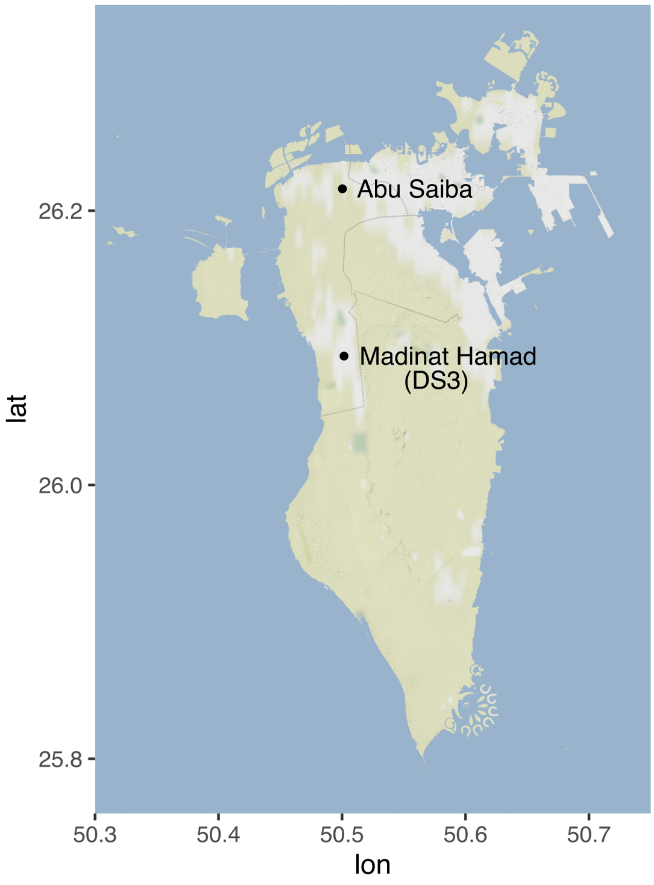


Figure S1 - Geographical locations of Abu Saiba and Madinat Hamad archaeological sites in Bahrain.

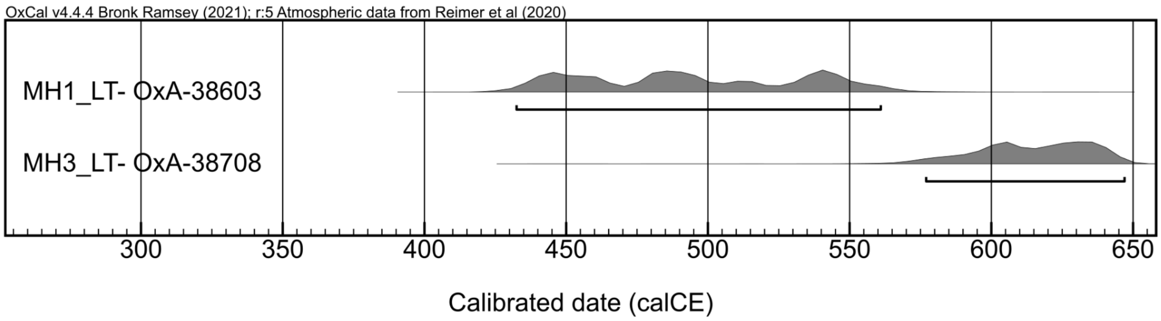


Figure S2 - Radiocarbon dating of two Bahrain samples, showing the density and 95.4% bars.

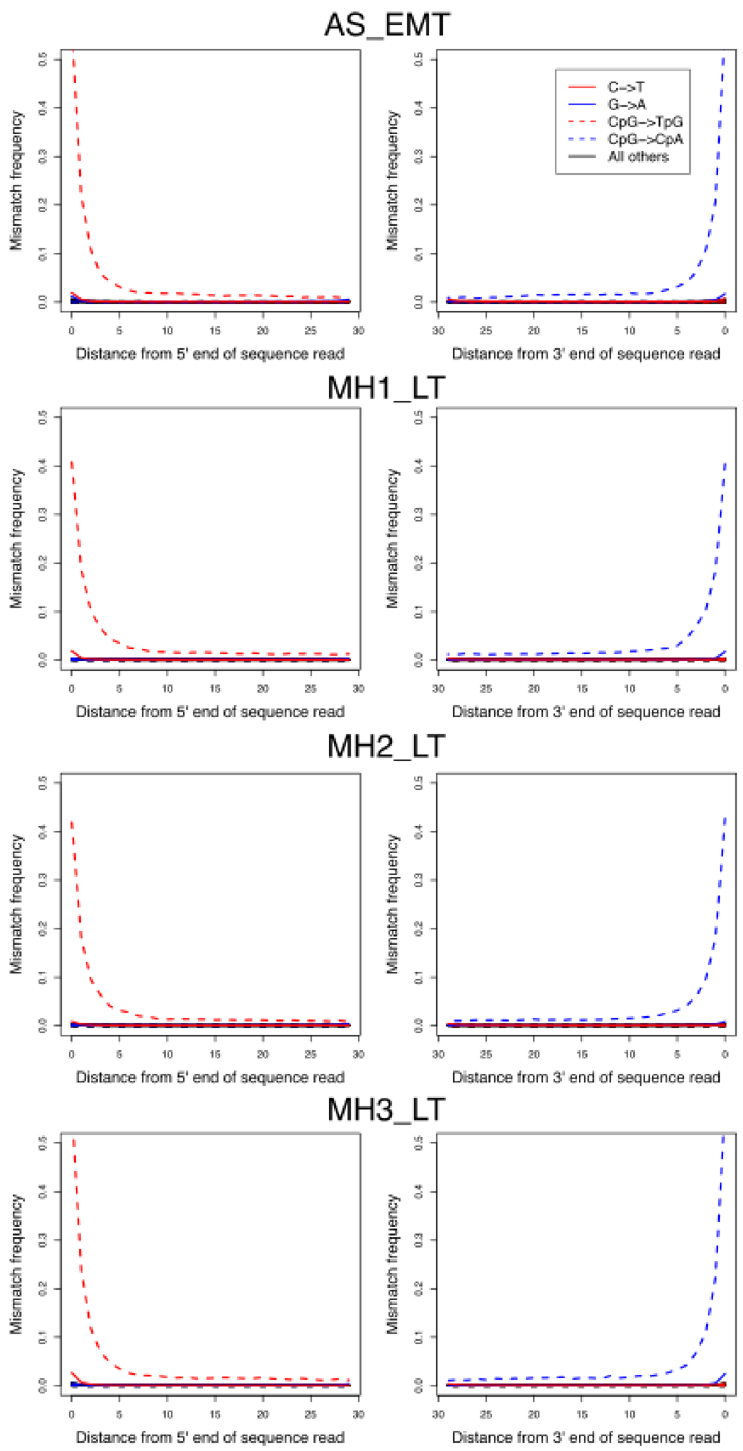


Figure S3 - Deamination patterns in the four ancient Bahrain samples. As expected in enzymatically treated samples, deamination is still observable at CpG sites.

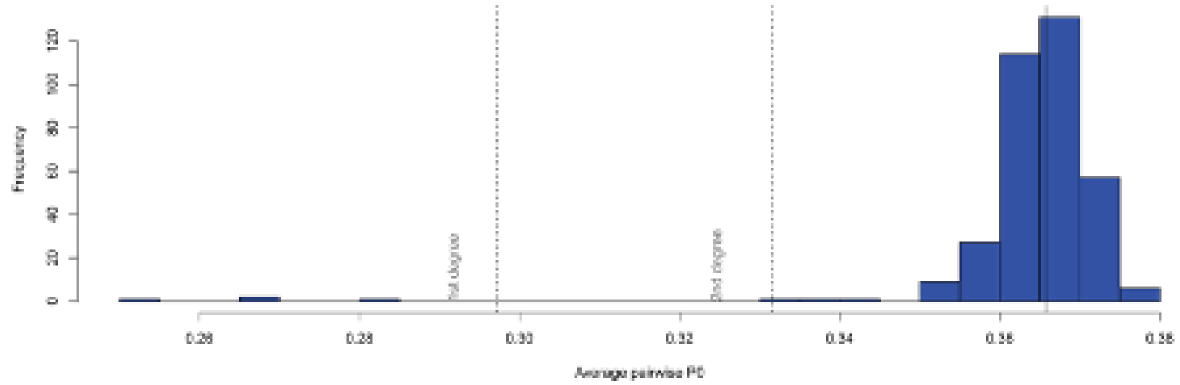


Figure S4 - Relatedness estimation using READ on Tylos-period Bahrain samples plus the following Samples/populations as controls: SFI-43 (Egyptian Mom), SFI-44 (SFI-43's son), Beirut_IA3, Beirut_Hellenistic, Beirut_IA2, Sidon_BA, Russia_Caucasus_Eneolithic, Russia_Caucasus_Eneolithic_sibling.I2056_sibling.I1722, Russia_Caucasus_Eneolithic_sibling.I2055_sibling.I2056.

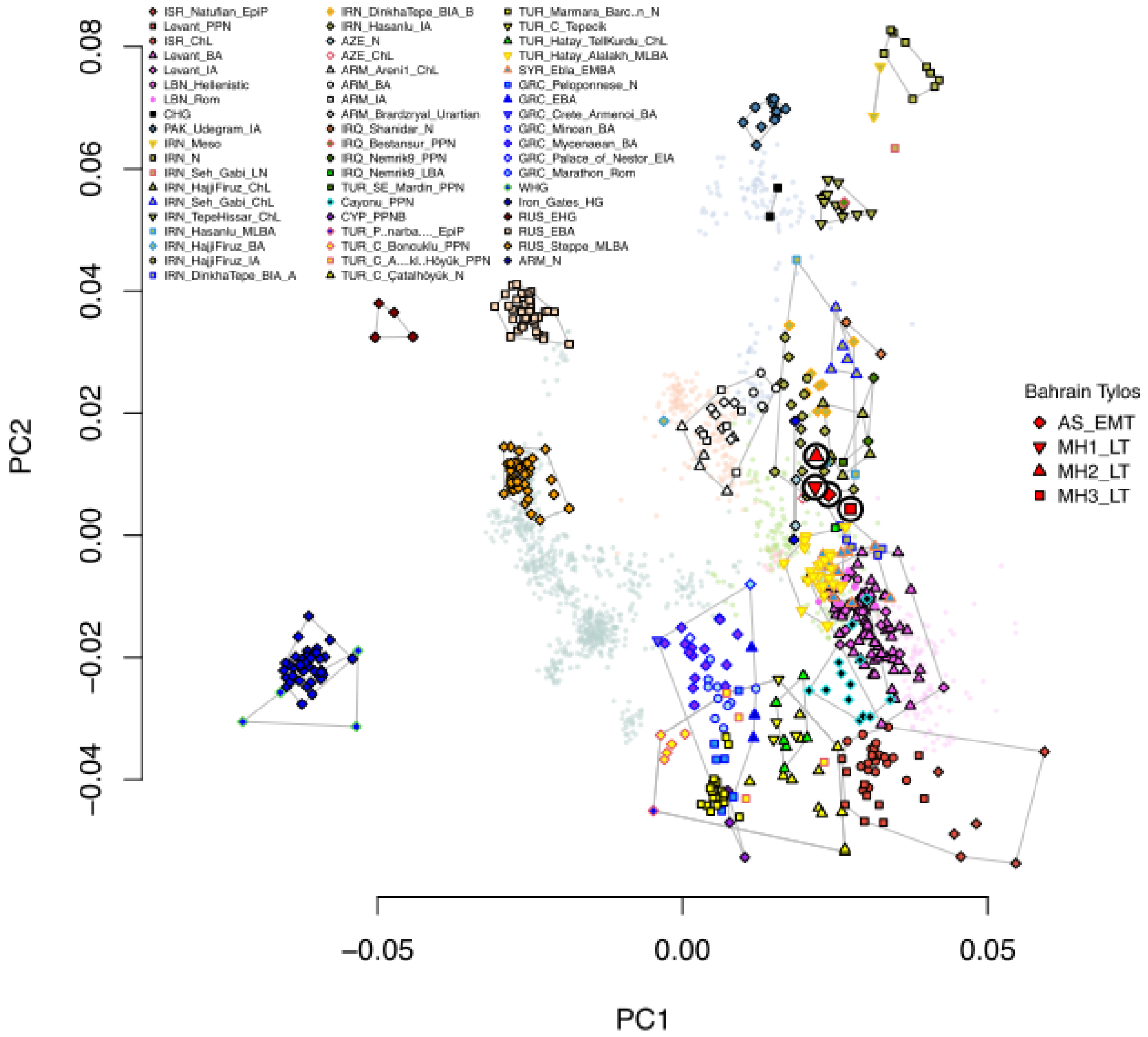


Figure S5 - Principal Component Analysis with 1,830 present-day and ancient Eurasians and 579,407 SNPs. Ancient samples are indicated with larger symbols as in the key.

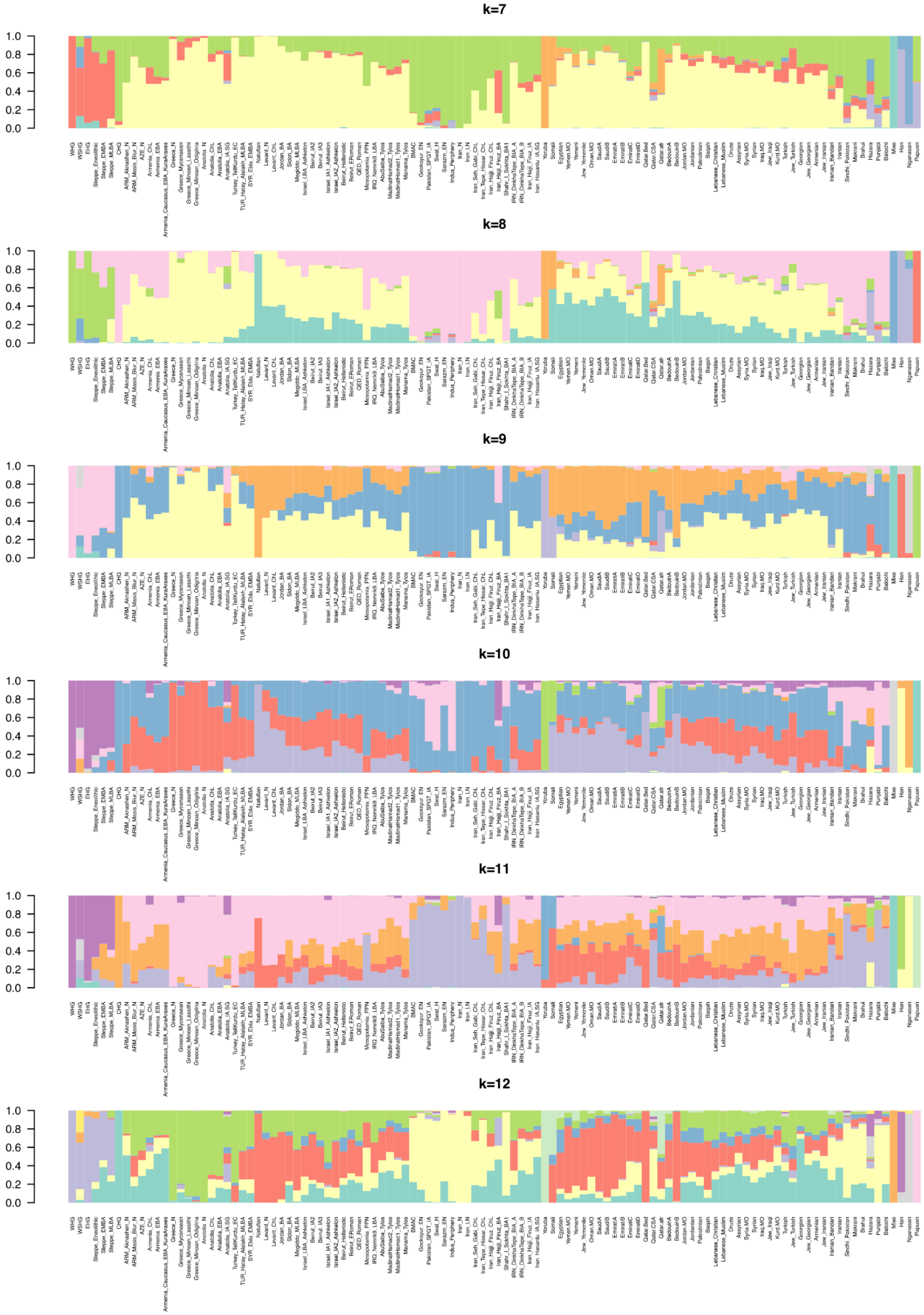


Figure S6 - Dystruct analysis k=7 to k=12.

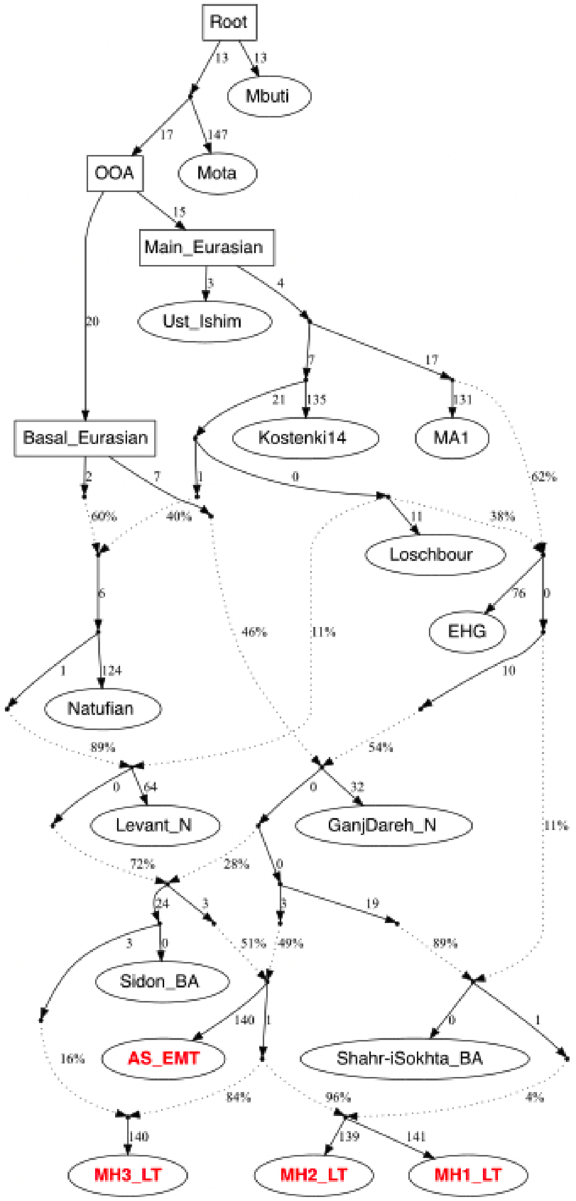


Figure S7 - Alternative model of Bahrain Tylos ancestry using qpGraph and IRN_Shahr-i Sokhta_BA instead of IRN_Hajji_Firuz_IA.

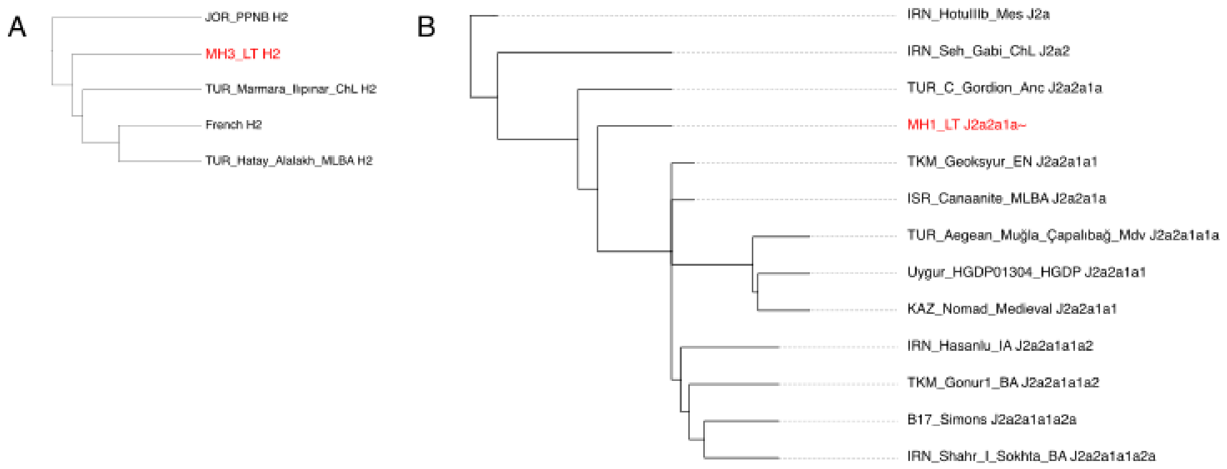


Figure S8 - pathPhynder placement of two Bahrain Late Tylos and other relevant ancient samples in a phylogeny of present-day and ancient Y-chromosome variation. A) H2 Y-chromosome lineages. B) J2a Y-chromosome lineages.

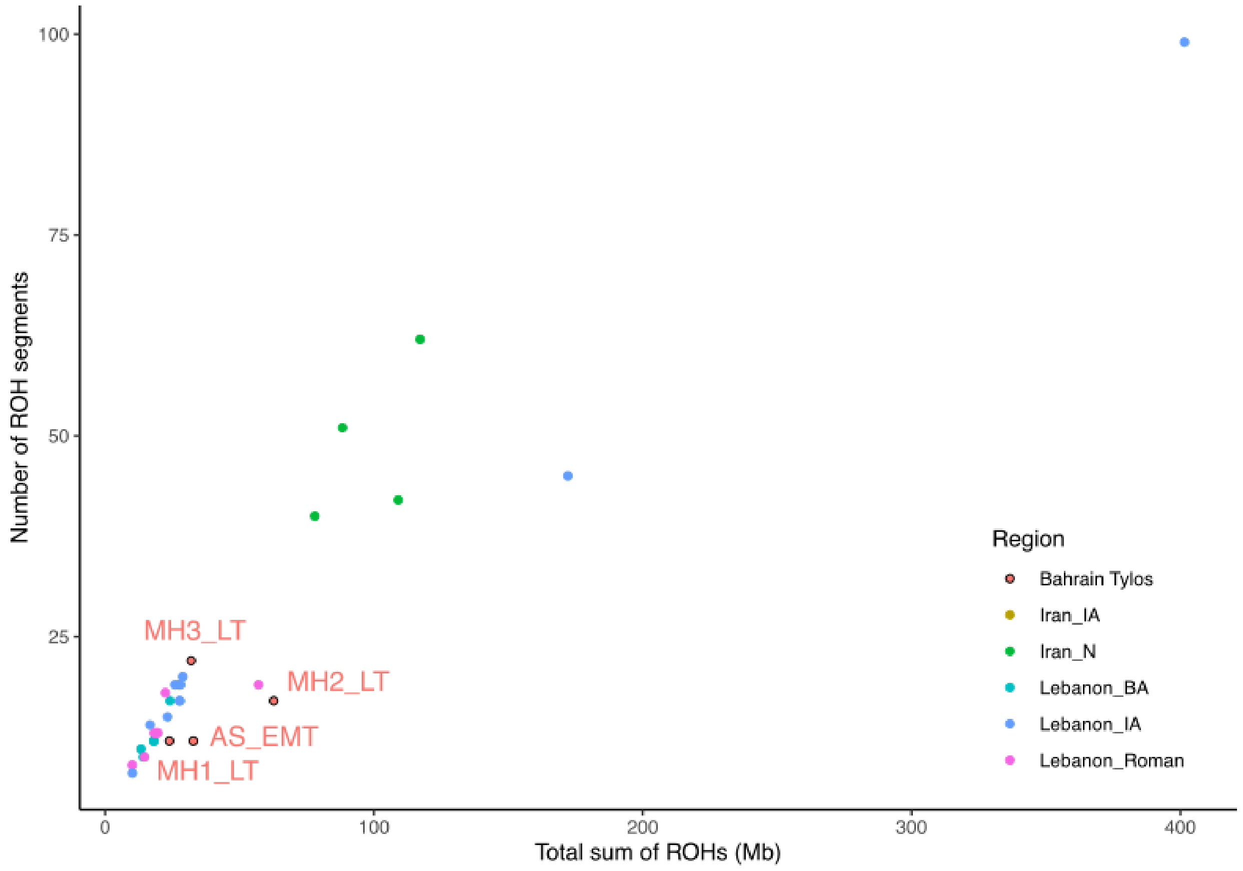


Figure S9 - Tylos-period Bahrain ROH distribution compared to ancient Levantine and Iranian samples.

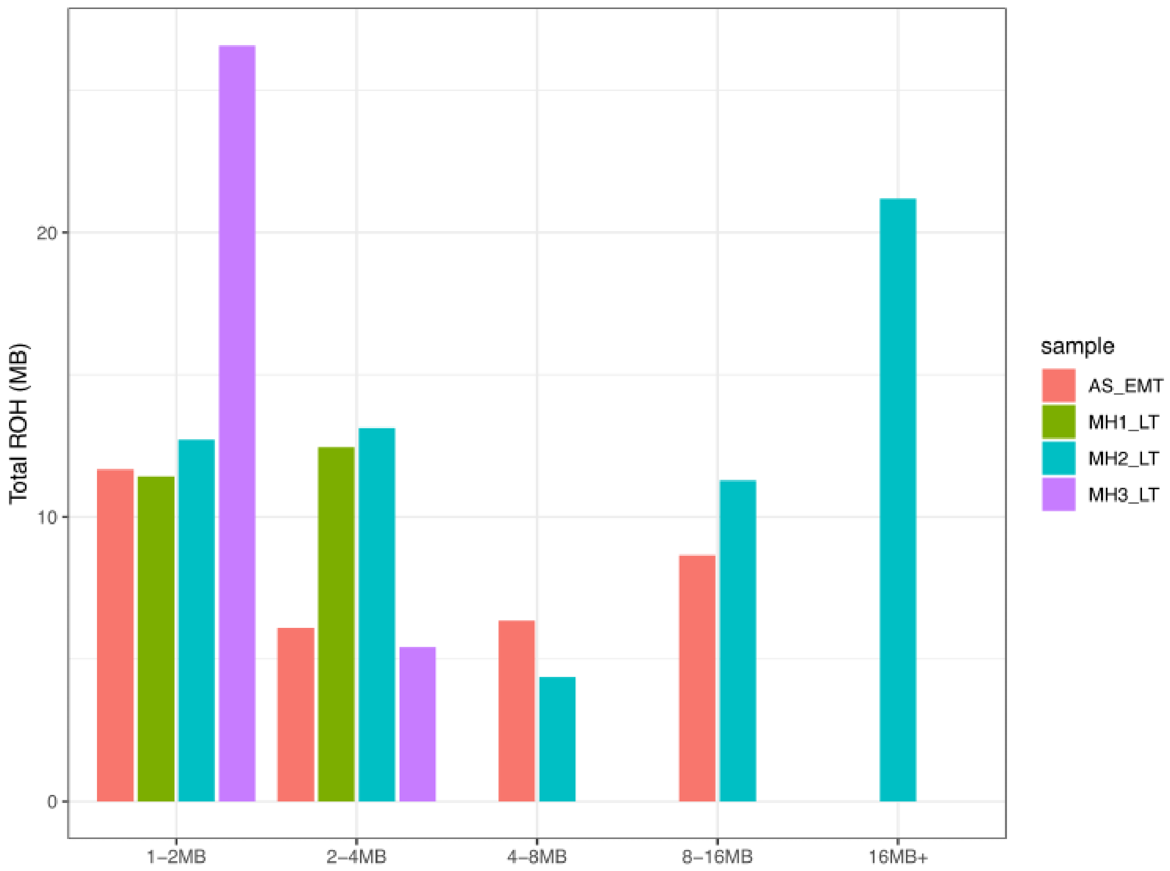


Figure S10 - ROH size distribution among Tylos-period Bahrainis.

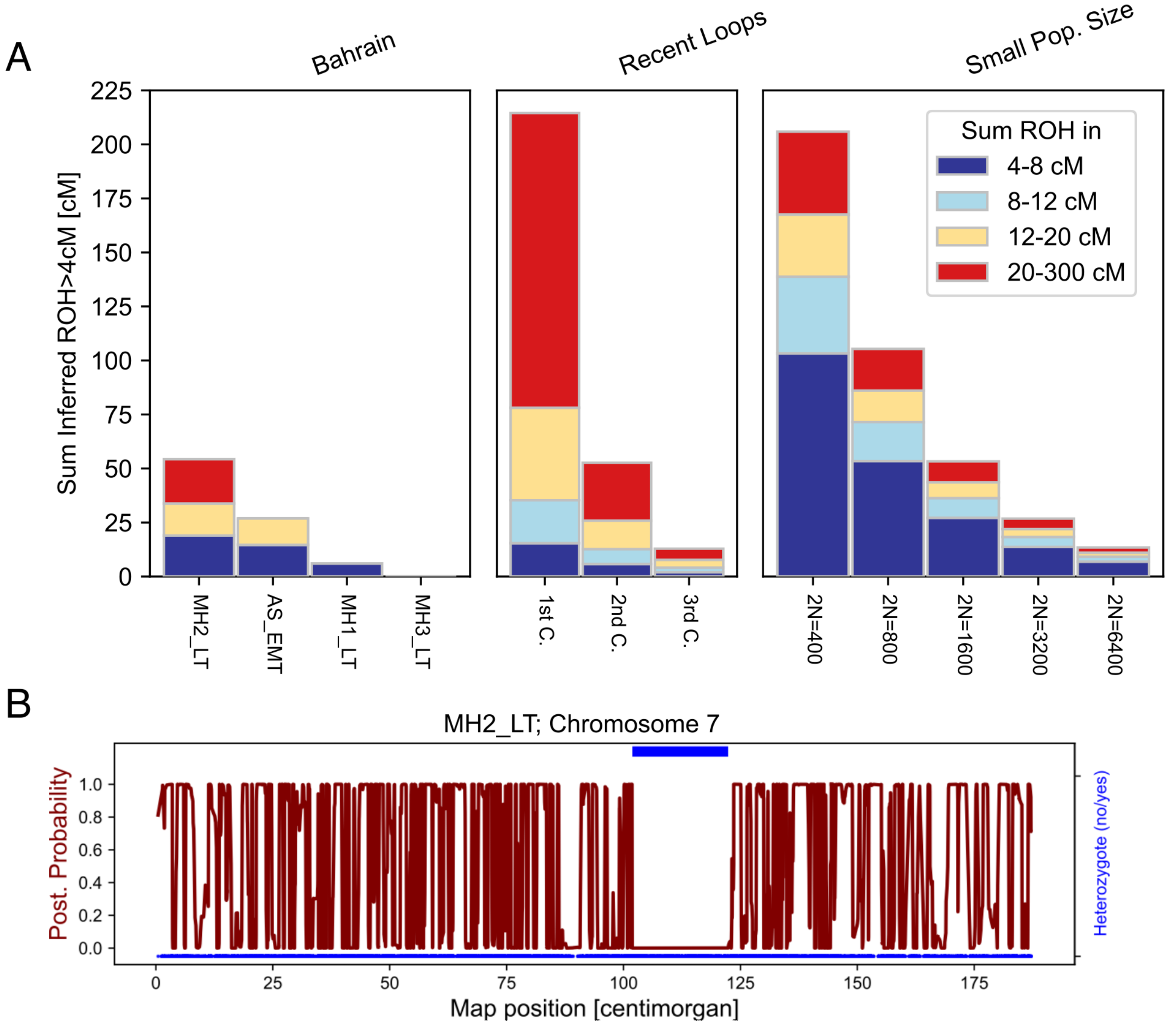


Figure S11 - A) Sum of inferred ROH>4cM in the four Tylos period samples from Bahrain and simulated proportions of ROH sizes in recent loops and small population sizes for comparison. B) ~20.47 cM segment of homozygosity found in the chromosome 7 of sample MH2_LT.

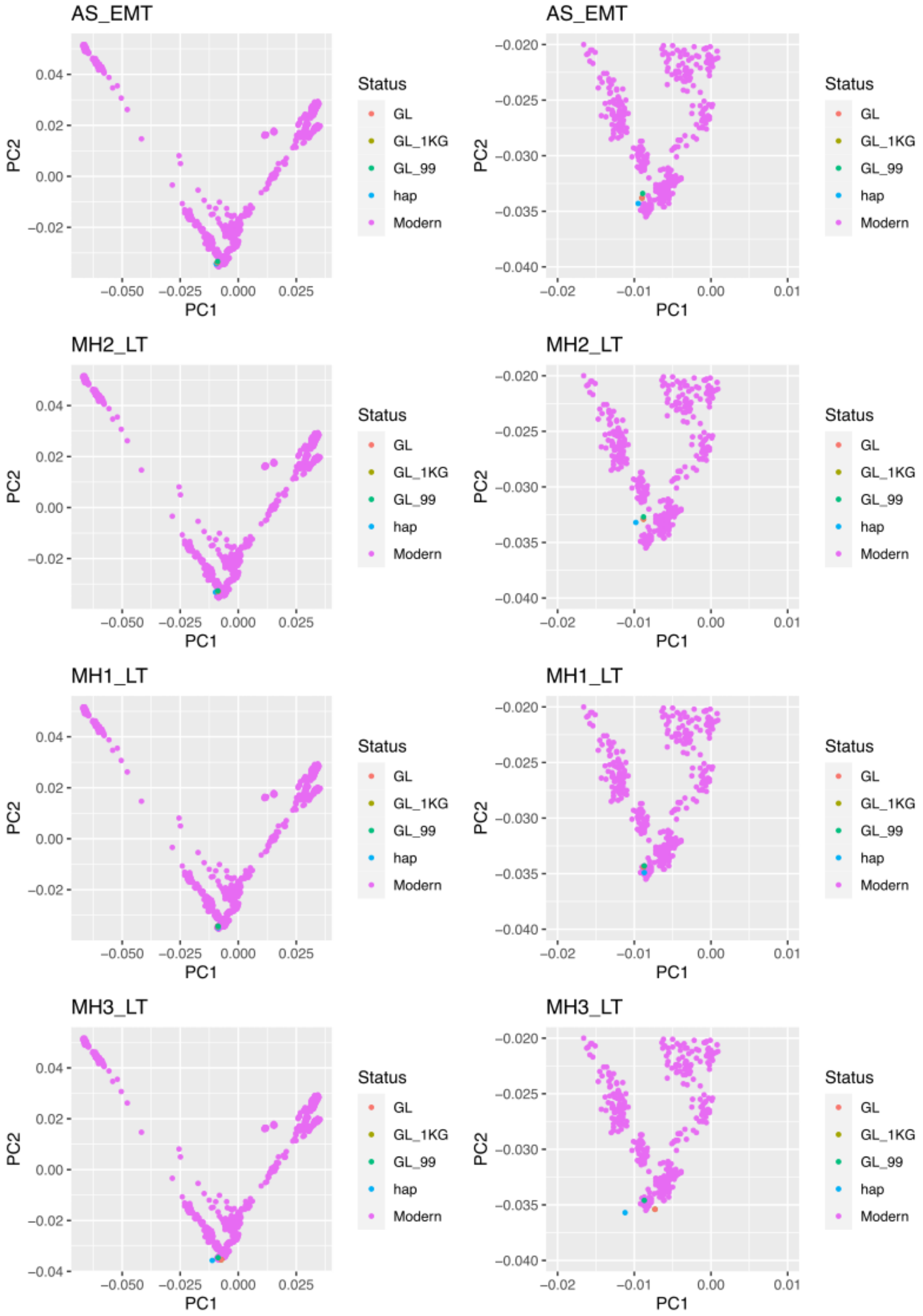


Figure S12 - Quality check of imputation. PCA calculated on the HGDP and projecting pseudo-haploid calls with different phased diploid calls.

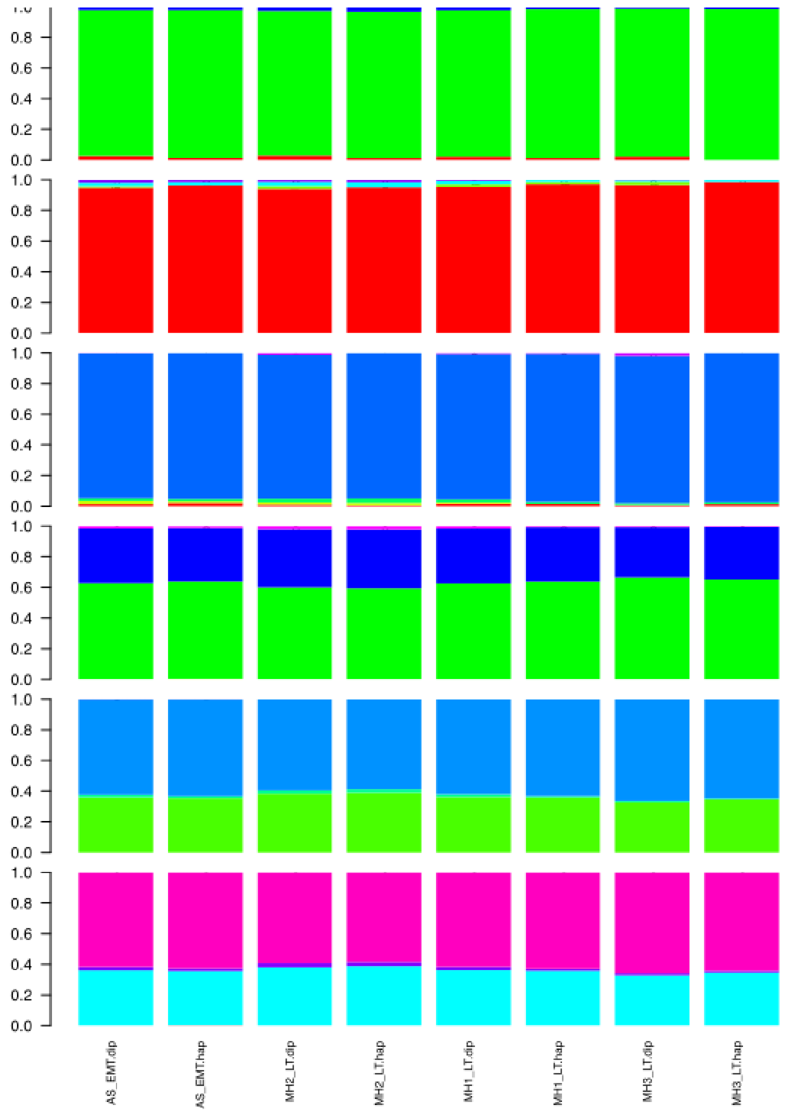


Figure S13 - Quality check of imputation. Admixture comparing pseudo-haploid calls with final imputed calls (GLIMPSE using HGDP + Almarri et al., 2021[16] as reference, setting GP <99 to missing and re-imputing the missing sites using the 1000G high coverage reference panel). K3 to K8, samples run with HGDP dataset.
